# “The Translesion Polymerase Pol Y1 is a Constitutive Component of the *B. subtilis* Replication Machinery”

**DOI:** 10.1101/2023.10.06.561215

**Authors:** McKayla E. Marrin, Michael R. Foster, Chloe M. Santana, Yoonhee Choi, Avtar S. Jassal, Sarah J. Rancic, Carolyn R. Greenwald, Madeline N. Drucker, Elizabeth S. Thrall

**Affiliations:** Department of Chemistry, Fordham University, Bronx, NY, 10458, USA

## Abstract

Unrepaired DNA damage encountered by the cellular replication machinery can stall DNA replication, ultimately leading to cell death. In the DNA damage tolerance pathway translesion synthesis (TLS), replication stalling is alleviated by the recruitment of specialized polymerases to synthesize short stretches of DNA near a lesion. Although TLS promotes cell survival, most TLS polymerases are low fidelity and must be tightly regulated to avoid harmful mutagenesis. The gram-negative bacterium *Escherichia coli* has served as the model organism for studies of the molecular mechanisms of bacterial TLS. However, it is poorly understood whether these same mechanisms apply to other bacteria. Here, we use in vivo single-molecule fluorescence microscopy to investigate the TLS polymerase Pol Y1 in the model gram-positive bacterium *Bacillus subtilis*. We find significant differences in the localization and dynamics of Pol Y1 in comparison to its *E. coli* homolog, Pol IV. Notably, Pol Y1 is constitutively enriched at or near sites of replication in the absence of DNA damage through interactions with the DnaN clamp; in contrast, Pol IV has been shown to be selectively enriched only upon replication stalling. These results suggest key differences in the roles and mechanisms of regulation of TLS polymerases across different bacterial species.

## Introduction

The DNA replication machinery, the replisome, copies genomic DNA with high accuracy and efficiency.(1) Yet replicative DNA polymerases can be blocked by the presence of unrepaired DNA damage on the template strand. In the DNA damage tolerance pathway translesion synthesis (TLS), conserved from bacteria to humans, specialized translesion polymerases copy damaged DNA that would otherwise stall replication.(2, 3) In contrast to high-fidelity replicative polymerases, most TLS polymerases are members of the error-prone Y-family.(4, 5) Thus, although TLS promotes cell survival under stress by avoiding the deleterious consequences of replication stalling, the activity of TLS polymerases must be tightly regulated to prevent harmful levels of mutagenesis.

The gram-negative bacterium *Escherichia coli* has served as the model system for TLS polymerase regulation in bacteria.(3) *E. coli* has three TLS polymerases (Pols II, IV, and V), two of which (Pols IV and V) are members of the Y-family. Pol II is a high-fidelity B-family polymerase; although capable of bypassing certain DNA lesions, its role in TLS is less clear.(3, 6, 7) All three *E. coli* TLS polymerases are transcriptionally upregulated by the SOS DNA damage response, a pathway that induces expression of several dozen DNA damage tolerance and repair factors in response to persistent single-stranded DNA (ssDNA).(8) Pols II and IV are expressed at moderate copy numbers, estimated as approximately 50 and 30 per cell, during normal growth.(9, 10) Upon SOS induction, they are upregulated approximately 10-fold. In contrast, Pol V is not expressed in the absence of induction and reaches levels of 15 – 60 copies per cell late in the SOS response.(9) Pols II and IV are single gene products, whereas Pol V contains UmuC and UmuD subunits produced from the *umuDC* operon.(3) Expression of Pol V is also regulated post-translationally in a pathway involving RecA that generates the active UmuD’_2_C form of the polymerase.

The activity of *E. coli* TLS polymerases also requires distinct protein-protein interactions. Pols IV and V must bind the ring-shaped replication processivity factor, the β clamp, to perform TLS(3); this interaction also stimulates DNA synthesis and lesion bypass by Pol II.(11) In addition, all three polymerases interact with another replisome component, single-stranded DNA-binding protein (SSB), via a conserved binding site at its C-terminal tail.(12, 13) Recent quantitative microscopy studies have revealed that these interactions play a role in the molecular regulation of TLS polymerase activity. In particular, Pol IV is not enriched at sites of replication during normal growth,(10, 14) but it is selectively recruited, primarily through the interaction with SSB, in response to DNA damage or other replication perturbations.(13–15) This selective enrichment likely helps to minimize mutagenesis in the absence of replication stress by limiting the access of Pol IV to the DNA template. Pol V appears to undergo a complex series of localization changes upon SOS induction; it was not observed to be enriched near sites of replication at any point after the induction of DNA damage.(16) These findings suggest a model in which spatial control is an important aspect of TLS polymerase regulation in *E. coli*.

Despite advances in elucidating the molecular mechanisms of TLS in *E. coli*, there has been relatively little work in other bacterial species. Thus, it remains unclear if these mechanisms are conserved. The model low-GC gram-positive bacterium *Bacillus subtilis* represents an attractive system for exploring TLS polymerase regulation in an evolutionarily distant species. Early studies in *B. subtilis* identified two putative Y-family TLS polymerases, Pol Y1 (encoded by the *yqjH* gene) and Pol Y2 (encoded by the *yqjW gene*).(17–19) These polymerases were determined to be homologous to *E. coli* Pol IV and the UmuC subunit of Pol V, respectively.(17, 20) However, there are clear differences in Pol Y1 and Pol Y2 regulation and activity in comparison to their *E. coli* homologs.

Like *E. coli* Pol IV, Pol Y1 is a single gene product; uncharacteristically for a TLS polymerase, however, it is not SOS-regulated.(18, 21) Pol Y1 contains a conserved clamp-binding motif (CBM), a pentapeptide sequence predicted to interact with the DnaN clamp (Figure 1A).(18, 22) The role of this clamp interaction in damage tolerance has not been determined, but a reduction in Pol Y1-mediated untargeted mutagenesis was observed upon mutation of the Pol Y1 CBM(18) and a Pol Y1-DnaN interaction was detected in a yeast two-hybrid assay.(23) The importance of other protein-protein interactions for Pol Y1 activity is unknown, although the same yeast two-hybrid assay detected an interaction with Pol I(23) and bioinformatics analysis suggested a possible interaction with RecA through a domain similar to the RecA N-terminal domain (RecA-NT) (Figure 1A).(24) Pol Y1 promotes survival upon treatment with various DNA damaging agents, including 4-nitroquinoline 1-oxide (4-NQO) and UV radiation.(17, 25) In contrast, Pol IV does not promote survival or mutagenesis upon UV exposure; instead, Pol V bypasses UV lesions in *E. coli*.(26) Finally, Pol Y1 has been implicated in responding to replication-transcription conflicts and appears to act in the transcription-coupled nucleotide excision repair (NER) pathway.(25) Similarly, there is evidence that Pol IV may play a role in transcription-coupled NER or TLS pathways in *E. coli*.(27, 28)

**Figure 1.**
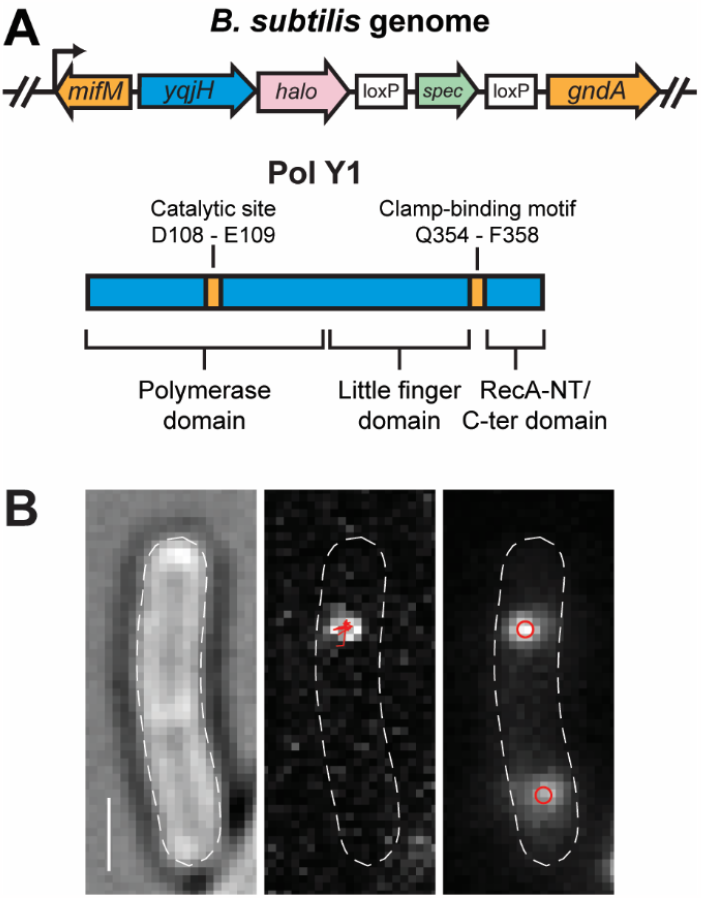
(A) Cartoons of (top) Pol Y1-Halo (*yqjH-halo*) fusion and context in *B. subtilis* genome and (bottom) Pol Y1 domain organization and key residues. (B) Representative micrographs recorded with 13.9 ms integration time. Left: transmitted white light micrograph of *B. subtilis* cell with overlaid cell outline and 1 μm scale bar. Middle: fluorescence micrograph of single Pol Y1-Halo-JFX_554_ molecule with overlaid trajectory. Right: fluorescence micrograph of DnaX-mYPet foci with overlaid centroids.

In this study, we use in vivo single-molecule fluorescence microscopy to quantify the location and dynamics of Pol Y1, both during normal growth and upon DNA damage with the drug 4-NQO. We observe static and mobile populations of Pol Y1 under both conditions, representing Pol Y1 molecules bound to DNA or diffusing in the cytoplasm, respectively. Notably, we find that the static population of Pol Y1 is moderately enriched near sites of replication during normal growth, in contrast to *E. coli* Pol IV. This enrichment does not require Pol Y1 catalytic activity, but it does require interactions with the DnaN clamp. Surprisingly, although Pol Y1 promotes survival in response to treatment with 4-NQO, there is little change in Pol Y1 localization and dynamics upon 4-NQO treatment, again in contrast to previous observations for Pol IV. Our results reveal significant differences in the activity of TLS polymerases across different bacterial species.

## Materials and Methods

### Bacterial Strain Construction

All bacterial strains were based on the wild-type (WT) *B. subtilis* background PY79.(29, 30) New bacterial strains were constructed by transformation of double-stranded DNA (dsDNA) fragments generated by polymerase chain reaction (PCR) and Gibson assembly(31) or of genomic DNA. Transformants were selected on antibiotic plates and validated by diagnostic PCR and Sanger DNA sequencing of the modified locus. All oligonucleotides and bacterial strains used in this study are listed in Tables S1 and S2. Detailed strain construction information is provided in the Supplementary Methods.

### Survival Assays

Glycerol stocks were streaked onto LB Lennox agar plates containing the appropriate antibiotics for strain selection (spectinomycin at 100 μg/mL or chloramphenicol at 5 μg/mL) and incubated overnight at 37 °C. Liquid cultures were prepared in LB Lennox media. The day before the experiment, a 2.5 mL culture of each strain was inoculated with a single colony and a second 2.5 mL culture was inoculated with a 1:100 dilution of the first culture. Both cultures were grown overnight for approximately 16 h at 22 ºC shaking at 225 rpm. The following morning, fresh cultures of 3 mL volume with an optical density at 600 nm (OD_600nm_) of 0.05 were prepared using the overnight culture with OD_600nm_ = 0.4 – 1.0. The fresh cultures were then grown at 37 ºC shaking at 225 rpm for approximately 3.5 h. LB Lennox agar plates were poured the day of the experiment and supplemented with a 1:1,000 dilution of a freshly prepared solution of 4-NQO in dimethylformamide (DMF), giving final concentrations of 0, 0.05, 0.075, or 0.1 μM 4-NQO. Serial 10-fold dilutions of the fresh cultures were prepared in LB Lennox media and 200 μL aliquots of appropriate dilutions were spread on the 4-NQO plates using glass beads. The plates were incubated at 37ºC overnight and colonies were enumerated the following morning. At least three independent replicates were performed for each experiment.

### Cell Culture and Sample Preparation for Microscopy

Glycerol stocks were streaked onto LB Lennox agar plates as described previously. Liquid cultures were grown in freshly prepared S7_50_-sorbitol minimal medium,(32) in which glutamate was used at 20 mM concentration and sorbitol at 0.4% concentration. The day before the experiment, a single colony was picked and used to inoculate a culture in 2.75 mL media, which was then added at 1:10 dilution into 2.25 mL media, yielding two cultures of 2.5 mL volume. Both cultures were grown overnight for approximately 16 h at 30 °C shaking at 225 rpm. The following morning, the culture with OD_600nm_ = 0.4 – 1.0 was used to inoculate a 25 mL culture for imaging to an initial OD_600nm_ = 0.005 – 0.02. Imaging cultures were grown at 37 °C shaking at 225 rpm until reaching early exponential phase, or OD_600nm_ ≈ 0.1 – 0.2.

When a culture was ready to harvest for microscopy, a 3 mL aliquot was pelleted by centrifugation at 3,500 × *g* for 2.5 min. All but 200 μL of the supernatant was aspirated and the cell pellet was resuspended to a final volume of 500 μL with fresh media. The sample was labeled by adding a 1:1,000 dilution of Janelia Fluor X 554 (JFX_554_)(33) HaloTag(34) ligand dissolved in dimethylsulfoxide (DMSO). The sample was then incubated for 15 min at 37 °C shaking at 225 rpm. The cell suspension was pelleted by centrifugation as before and the supernatant was removed. The cells were washed by adding 1 mL of fresh media, resuspending the pellet, and centrifuging again. Most of the supernatant was removed and the pellet was resuspended in the remaining few μL of liquid. A small volume (< 1 μL) of the concentrated cell suspension was deposited on an agarose pad and sandwiched between two coverslips for imaging.

Agarose pads were prepared by melting GTG agarose (NuSieve) at 3% concentration in S7_50_-sorbitol media at 65 °C for 15 – 30 min. Coverslips were cleaned by rinsing with ethanol and deionized (DI) H_2_O and dried in a nitrogen gas stream. To cast an agarose pad, 500 μL of molten agarose was deposited between two coverslips and left to cool for 20 – 30 min until solid. Coverslips in contact with the microscope objective were cleaned by 30 min cycles of sonication alternating between alcohol and 1 M KOH twice each, stored in DI H_2_O, and rinsed with DI H_2_O before use.

### Sample Treatment for Microscopy

For 4-NQO treatment of imaging cultures, a freshly prepared 4-NQO solution in DMF was diluted 1:1,000 into an early exponential phase (OD_600nm_ ≈ 0.1 – 0.2) culture. The treated culture was grown shaking at 37 °C and 225 rpm for an additional 1 or 2 hours. The effect of the treatment on cell growth was determined by measuring the density of colony-forming units (CFUs) per mL of culture before and after treatment. In brief, an aliquot of culture was diluted serially in 0.9% NaCl and a 200 μL volume was deposited on plain LB Lennox agar plates and spread with glass beads. Plates were incubated overnight at 37 °C, colonies were enumerated, and the number of CFUs/mL was calculated.

For fixed cell imaging experiments, a 4.5 mL aliquot of culture was harvested in early exponential phase (OD_600nm_ ≈ 0.1 – 0.2) and concentrated by two rounds of centrifugation at 3,500 × *g* for 2.5 min. The resulting cell pellet was resuspended at 3× concentration in a 2.5% solution of formaldehyde in phosphate buffered saline (PBS) and incubated with gentle shaking at room temperature for 45 min. Cells were then pelleted by centrifugation as before and resuspended in 500 μL of fresh S7_50_-sorbitol media, then labeled with JFX_554_ HaloTag ligand and prepared for microscopy as described previously.

### Microscopy

Fluorescence imaging was performed on a Nikon Ti2-E microscope equipped with a Nikon CFI Apo 100x/1.49 NA total internal reflection fluorescence (TIRF) objective lens and a Hamamatsu ImageEM C9100-23BKIT EMCCD camera. An additional internal magnification of 1.5× resulted in a pixel size of approximately 106 nm, determined by measuring a calibration grid (Thorlabs R1L3S2P). Fluorescence excitation was provided by 514 nm (150 mW) and 561 nm (200 mW) Coherent Sapphire lasers; laser powers were adjusted using neutral density filters. The individual laser beams were first expanded with a telescope and then passed through excitation filters (Chroma ZET514/10x and ZET561/10x) before being combined with a mirror and dichroic filter (Chroma ZT514rdc-UF2). After passing through a second dichroic filter (Chroma ZT405rdc-UF2) for an unused laser line, the beams were then expanded with a second telescope and focused to the objective back focal plane (BFP) using a 400 mm focal length lens (Thorlabs AC508-400-A) mounted on a micrometer stage. Highly inclined thin illumination,(35) or near-TIRF, excitation was achieved by translating the lens to move the focused beam away from the center of the objective BFP. The microscope contained a TIRF filter cube (Chroma) with multi-band dichroic and emission filters (Chroma ZT405/514/561rpc-UF2 and ZET442/514/561m) and a longpass filter (Chroma ET525lp) to remove the unnecessary short-wavelength passband. Brightfield images of cells were recorded with white light transillumination. The microscope was equipped with a computer-controlled translation stage (Mad City Labs MicroStage). Computer-controlled shutters (Vincent Uniblitz VS14ST0 and VMM-D4 driver) were used to automate laser excitation.

Movies were recorded with either a short integration time (13.9 ms) to enable tracking of all labeled molecules or a long integration time (250 ms) for selective imaging of static molecules.(14, 36) For single Pol Y1-Halo-JFX_554_ molecules, 561 nm laser powers of approximately 15 and 5 W/cm^2^ were used for short- and long-exposure imaging, respectively. For DnaX-mYPet foci, 514 nm laser powers of approximately 1 and 0.1 W/cm^2^ were used for short- and long-exposure imaging, respectively. All movies commenced with 100 frames of 514 nm excitation before either continuous or stroboscopic 561 nm excitation.

To check for any apparent offset between the 514 nm and 561 nm channels due to chromatic aberrations in the microscope optics, we imaged 0.1 μm diameter TetraSpeck beads (Thermo Fisher Scientific #T7279), used u-track to determine the bead centroids, and compared the apparent centroid positions under 514 nm and 561 nm excitation. We found an offset of approximately 11 nm between the two channels, indicating that chromatic effects are negligible.

### Image Analysis

Automated image analysis was performed using the MATLAB-based packages MicrobeTracker (v. 0.937)(37) and u-track (v. 2.1.1),(38, 39) in addition to custom MATLAB code. As the initial step, cell segmentation was performed on brightfield images using MicrobeTracker to generate cell outlines. Spot detection was performed within each individual cell outline using the u-track point source detection algorithm, which fit spots to symmetrical two-dimensional (2D) Gaussian approximations of the point spread function (PSF); fit parameters consisted of the *x* and *y* centroid positions, the Gaussian width, the Gaussian amplitude, and the background offset. Detected spots were then linked in trajectories using the u-track tracking algorithm. Nearest neighbor distance was not used to expand the search radius. Analysis settings for specific movie types are described below; if not otherwise noted, default spot detection and tracking parameters were used. A small number of cells were manually excluded from analysis if they contained an unusually high fluorescent background.

For short-exposure Pol Y1-Halo imaging, the significance threshold *α* = 10^−5^ was used. Gaps of one frame duration were allowed in trajectories to account for missed localizations due to fluorophore blinking or failed detection. Tracks were required to contain at least two localizations, although tracks containing fewer than five localizations were excluded from subsequent analysis. The Brownian search radius multiplier factor was set to 5.

For long-exposure Pol Y1-Halo imaging, the *α* = 10^−6^ threshold was used and no gaps were allowed in trajectories. The upper bound on the Brownian search radius was set to 3 pixels. After spot detection and tracking, static molecules were identified by requiring the average of the PSF width for all localizations within a trajectory to be in the range of 0.703 – 2.145 pixels (74.5 – 227.4 nm). These values were taken as the mean value ± two standard deviations of a Gaussian fit to the distribution of PSF widths observed in fixed cells (Figure S1C); this requirement minimizes spurious detection of mobile molecules, which give rise to spots with broader PSFs in long-exposure imaging.(14, 36)

For both exposure times, DnaX-mYPet foci were analyzed by generating an average projection of the first 20 frames of 514 nm excitation. Spot detection was performed on the resulting average image as described above, using a significance threshold of *α* = 10^−5^. To remove a small number of false positive spots, the Gaussian background value was required to be above the camera offset level of 2,065 counts.

### Imaging Dataset

In general, experiments were performed on at least two different days with at least three independent replicates, defined as separate imaging cultures. Some control experiments, however, were performed with only two replicates. In all cases, individual replicates were checked for consistency. Table S3 lists the number of imaging days, replicates, cells, and tracks or foci for all imaging data.

### Data Analysis

#### Diffusion Coefficient Analysis

Apparent 2D diffusion coefficients (*D*^*^) were calculated from the mean squared displacement (MSD) of short-exposure trajectories with at least five localizations as:

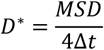

where Δ*t* is the time interval between subsequent localizations. The MSD is given by:

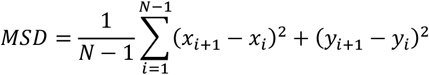

where *x*_i_ and *y*_i_ are the track coordinates in the *i*^th^ frame.

The resulting distributions for *D*^*^ revealed the presence of more than one diffusing species. These probability distributions were fit to a model for three diffusing species, which gave the best agreement with the experimental results. For trajectories containing exactly four steps, the following analytical expression describes the probability distribution:

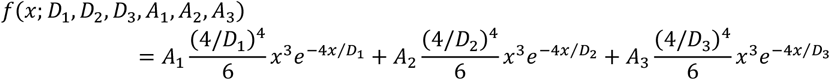

where *A*_n_ is the fraction of molecules in the *n*^th^ species and *D*_n_ represents the diffusion coefficient of that species.(15, 40) Fitting was performed subject to the normalization constraint:

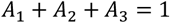

Longer trajectories were truncated to retain only the first four steps before use in this analysis.

To differentiate between static and mobile Pol Y1 molecules using a single *D*^*^ threshold value, we generated a distribution of observed *D*^*^ values in fixed cells (Figure S1B). Molecules were considered static if they had a *D*^*^ value less than the mean of a Gaussian fit to this distribution plus two standard deviations, corresponding to a cutoff of *D*^*^ < 0.14 μm^2^/s.

#### Cellular Localization Analysis

To allow comparison of Pol Y1-Halo and DnaX-mYPet localization across cells of different sizes and orientations, cell coordinates were rotated and normalized so that *x* and *y* coordinates spanned values from 0 – 1 along the long and short cellular axes, respectively.(14) First, one cell pole was set to a coordinate of (0,0) by subtracting the appropriate offset values from the cell outline *x* and *y* coordinates. The cell outline was then rotated by the angle between the two cell poles and the *x*-axis to align the cellular long axis with the *x*-axis. The cell outline *x* and *y* coordinates were divided by the cell length and width respectively and offset again such that the coordinates ranged from 0 – 1 along both cellular axes. The same transformations were then applied to the coordinates of Pol Y1 trajectories or DnaX foci to convert them to normalized coordinates. Finally, distributions of normalized *x* and *y* coordinates were generated. For Pol Y1, the mean *x* and *y* positions for each trajectory were compiled.

#### Radial Distribution Function Analysis

Intracell colocalization between individual Pol Y1-Halo molecules and DnaX-mYPet foci was quantified using radial distribution function analysis.(14, 41, 42) First, the average distance between each Pol Y1 trajectory and the closest DnaX focus was determined. Next, the same number of Pol Y1 localizations were randomly simulated within the same cell outline and the corresponding Pol Y1-DnaX distances were determined. Distributions of experimental and random Pol Y1-DnaX distances were generated by repeating this analysis for all cells in the dataset. If Pol Y1 were randomly localized relative to DnaX, these distance distributions would be identical. Finally, the radial distribution function, *g*(*r*), was generated by normalizing the experimental distribution by the random distribution. Random localization is indicated by *g*(*r*) ≈ 1 across all Pol Y1-DnaX distances *r*, whereas enrichment of Pol Y1 near DnaX is indicated by *g*(*r*) > 1 at short distances. To minimize variability, this procedure was repeated 100 times using 100 different random distance distributions and the resulting 100 *g*(*r*) curves were averaged. The standard error of the mean (S.E.M.) for these 100 *g*(*r*) curves are reported in Table S4. As a confirmation that the sample size was adequate, an independent random Pol Y1-DnaX distance distribution was generated and 100 random *g*(*r*) curves were calculated and averaged in the same manner. Deviations in the resulting random *g*(*r*) curve from 1 are due to the finite sample size. This radial distribution function approach accounts for spurious Pol Y1-DnaX colocalization due to cellular confinement; it also corrects for differences in cell size or number of DnaX foci across different treatment conditions.

#### Binding Lifetime Analysis

The Pol Y1 binding lifetime was measured by selectively imaging static molecules using long 250 ms integration times. For each static trajectory, the apparent binding lifetime was taken as the trajectory duration. Because this apparent binding lifetime includes both true dissociation events and events where the JFX_554_ fluorophore photobleaches, observed apparent binding lifetimes (*τ*_app_) were corrected by the photobleaching lifetime of JFX_554_ (*τ*_bleach_) to give a photobleaching corrected binding lifetime (*τ*_bound_):(36)

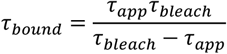

To determine *τ*_bleach_, JFX_554_-labeled Pol Y1 was imaged in fixed cells, where dissociation does not occur. The lifetimes *τ*_app_ and *τ*_bleach_ were determined from single exponential fits to the distributions of measured trajectory durations. Because these durations are not well fit by a single exponential and include both short and long timescale decays, we excluded tracks less than five frames in duration from the fits to capture the long timescale behavior.

#### Statistical Analysis

Distributions of the number of DnaX-mYPet foci per cell were compared using a two-sided Wilcoxon rank sum test, with a threshold of *p* < 0.05 for statistical significance.

## Results

### Creation and validation of a functional Pol Y1 fusion

To visualize single copies of Pol Y1 in live *B. subtilis* cells, we created a C-terminal fusion of the self-labeling HaloTag(34) to the endogenous copy of the *yqjH* gene (Figure 1A). By treating cells with a low (2.5 nM) concentration of a Janelia Fluor X 554 (JFX_554_) HaloTag ligand,(33) we were able to sparsely label Pol Y1 in live cells (Figure 1B, left panel) with the bright and photostable fluorophore JFX_554_, enabling detection and tracking of single Pol Y1 molecules (Figure 1B, middle panel). In addition, we introduced an orthogonal fusion of the yellow fluorescent protein mYPet(43) to the clamp-loader component DnaX; this DnaX-mYPet fusion forms bright and distinct foci, allowing simultaneous visualization of sites of DNA replication in the cell (Figure 1B, right panel).(44–47) To visualize the entire population of Pol Y1 molecules, we imaged cells with a short 13.9 ms exposure time. Alternatively, to resolve immobile Pol Y1 molecules selectively, we used a longer 250 ms exposure time; long exposure times blur out the fluorescence signal from rapidly-moving molecules such that only immobile molecules are detected.(14, 36)

High levels of background fluorescence have been observed in *B. subtilis* cells, potentially leading to detection of false positives in sensitive single-molecule fluorescence microscopy.(48) To address this possibility, we imaged a strain bearing only the DnaX-mYPet fusion labeled with the same 2.5 nM JFX_554_ dye concentration and analyzed the resulting false positive signal. Results are shown in the Supporting Information for localization (Figures S2A and B) and diffusion (Figures S5A – D) measurements. In all cases, we observed almost no false positive signal in comparison to the true Pol Y1-Halo-JFX_554_ signal.

We assessed the functionality of the Pol Y1-Halo fusion by assaying cell sensitivity to the DNA damaging agent 4-NQO, which generates DNA lesions on guanine and adenine bases, including quinoline adducts to the guanine *N*^2^ position.(49–51) Consistent with a previous report,(25) we find that cells lacking Pol Y1, but not Pol Y2, are sensitized to 4-NQO treatment (Figure 2A). Cells bearing the Pol Y1-Halo fusion are indistinguishable from WT, as are cells containing both the Pol Y1-Halo and DnaX-mYPet fusions (Figure 2B). We also constructed C-terminal Pol Y1 fusions to mYPet and to the photoactivatable fluorescent protein Dendra2,(52, 53) the latter of which was slightly sensitized to 4-NQO treatment relative to WT (Figure S1A).

**Figure 2.**
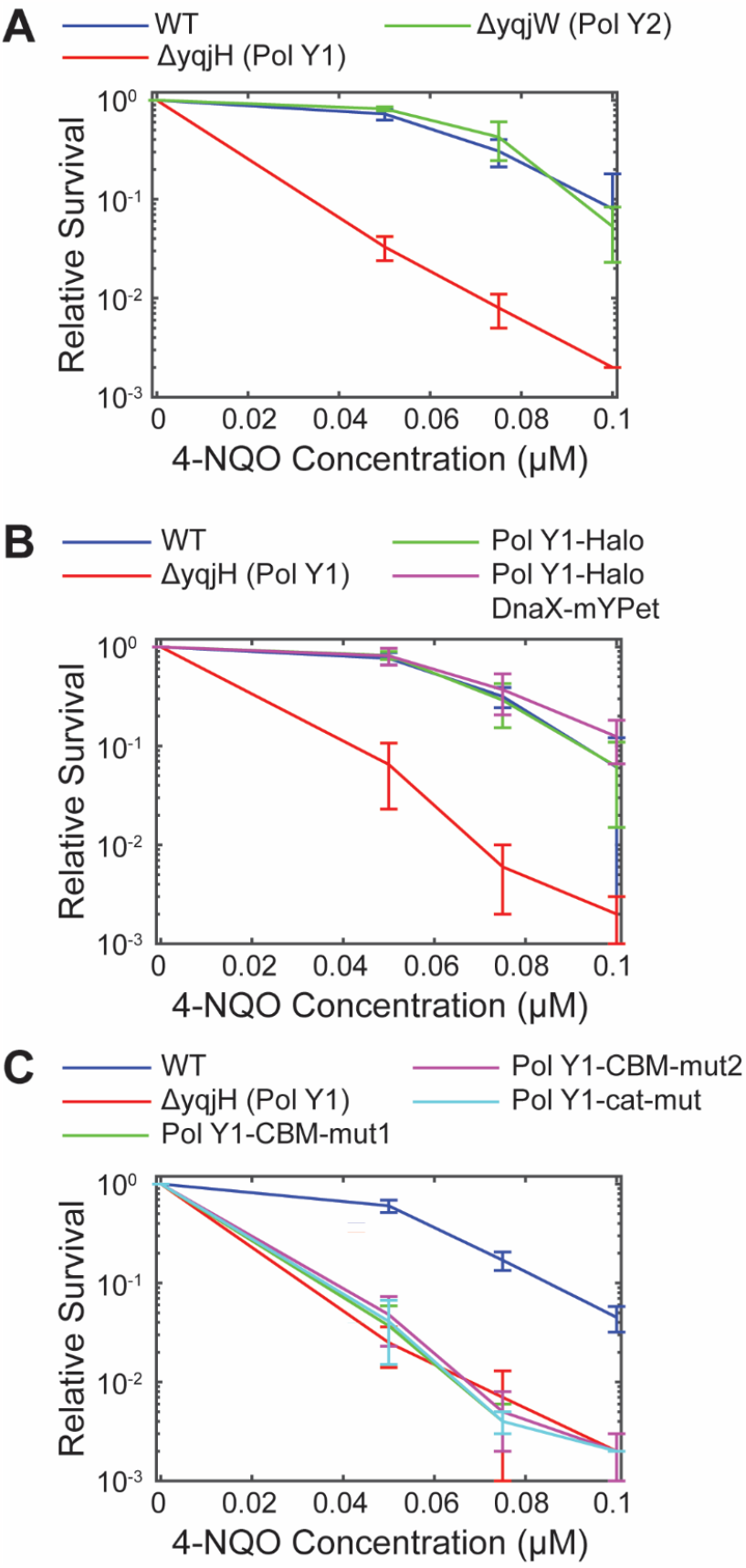
Relative survival of *B. subtilis* strains treated with different concentrations of 4-NQO. (A) WT Pol Y1, Pol Y1 knockout, and Pol Y2 knockout strains. (B) WT Pol Y1, Pol Y1 knockout, Pol Y1-Halo fusion, and Pol Y1-Halo fusion plus DnaX-mYPet fusion strains. (C) WT Pol Y1, Pol Y1 knockout, Pol Y1 clamp-binding mutant (Pol Y1-CBM-mut1 and Pol Y1-CBM-mut2), and Pol Y1 catalytically inactive mutant (Pol Y1-cat-mut) strains. Error bars show standard deviation of at least three replicates.

### Pol Y1 is moderately enriched near sites of replication during normal growth

In *E. coli*, TLS polymerases are largely excluded from the replication fork in the absence of DNA damage or other replication perturbations.(10, 14, 15) First, we asked whether the same was true for Pol Y1 by comparing the average cellular localization of Pol Y1 and DnaX. We found that DnaX foci were primarily localized at the quarter and three-quarter positions along the long cell axis and at midcell along the short cell axis (Figures 3A and 3B). This localization is consistent with previous reports of replication fork positioning in *B. subtilis*.(45, 46) Cells typically contained between one and two DnaX foci (mean ± S.E.M.: 1.67 ± 0.03). Pol Y1 localization was like that of DnaX, with strong enrichment at the quarter and three-quarter cell long axis positions and at midcell along the short axis (Figure 3C). We observed similar localization using a longer 250 ms integration time to resolve static Pol Y1 molecules selectively, but with greater Pol Y1 localization at the midcell position along the long cell axis (Figures S2C and E). Although these results are aggregated across unsynchronized cell populations, they suggest that Pol Y1 is enriched at or near sites of replication.

**Figure 3.**
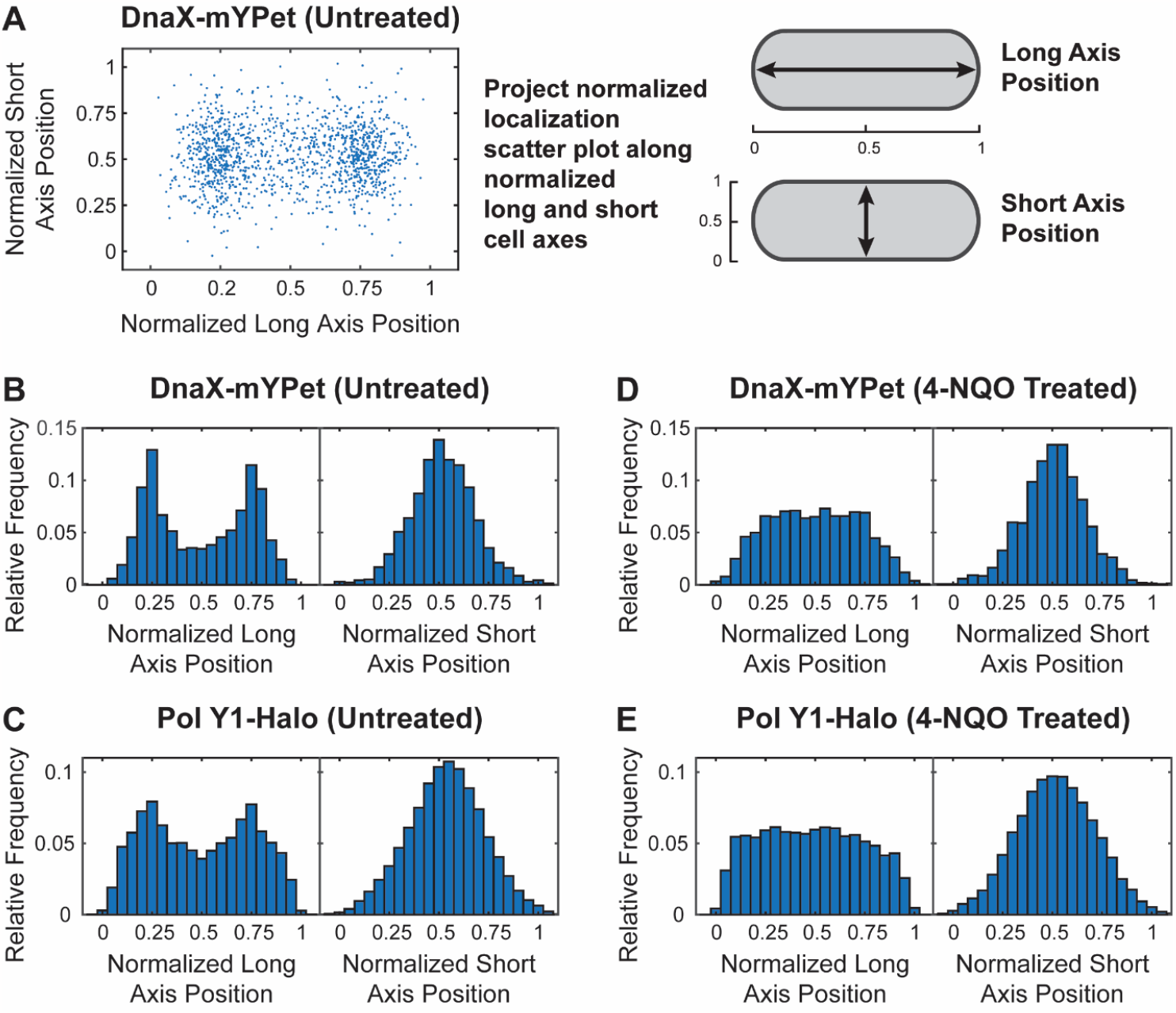
Cellular localization of DnaX-mYPet and Pol Y1-Halo. (A) Scatter plot of normalized positions of DnaX foci in untreated cells and cartoon of long and short cell axis projections. Long and short cell axis projections of (B, D) DnaX foci and (C, E) Pol Y1 trajectories in untreated cells and cells treated with 10 μM 4-NQO for 1 h, respectively.

To assess intracellular colocalization more rigorously, we used radial distribution function analysis. In this approach, the radial distribution function, *g*(*r*), represents the fold-enrichment of Pol Y1 molecules as a function of distance *r* from DnaX relative to random chance (Figure S4A). A *g*(*r*) value of one indicates no enrichment relative to chance, whereas higher values indicate greater colocalization.(14, 41) We observed moderate enrichment of Pol Y1 near DnaX, with a maximum *g*(*r*) ≈ 2.84 (Figure 4A); again, the enrichment was similar when using longer 250 ms integration times (Figure S4B). (See Table S4 for quantification of variability in all *g*(*r*) measurements.) In contrast, the *E. coli* homolog of Pol Y1, Pol IV, is only weakly enriched at sites of replication in the absence of DNA damage or other replication perturbations, with a maximum *g*(*r*) ≈ 1.5.(13–15) This difference in degree of replication fork enrichment between Pol Y1 and Pol IV suggests differences in TLS polymerase regulation and activity between *B. subtilis* and *E. coli*.

**Figure 4.**
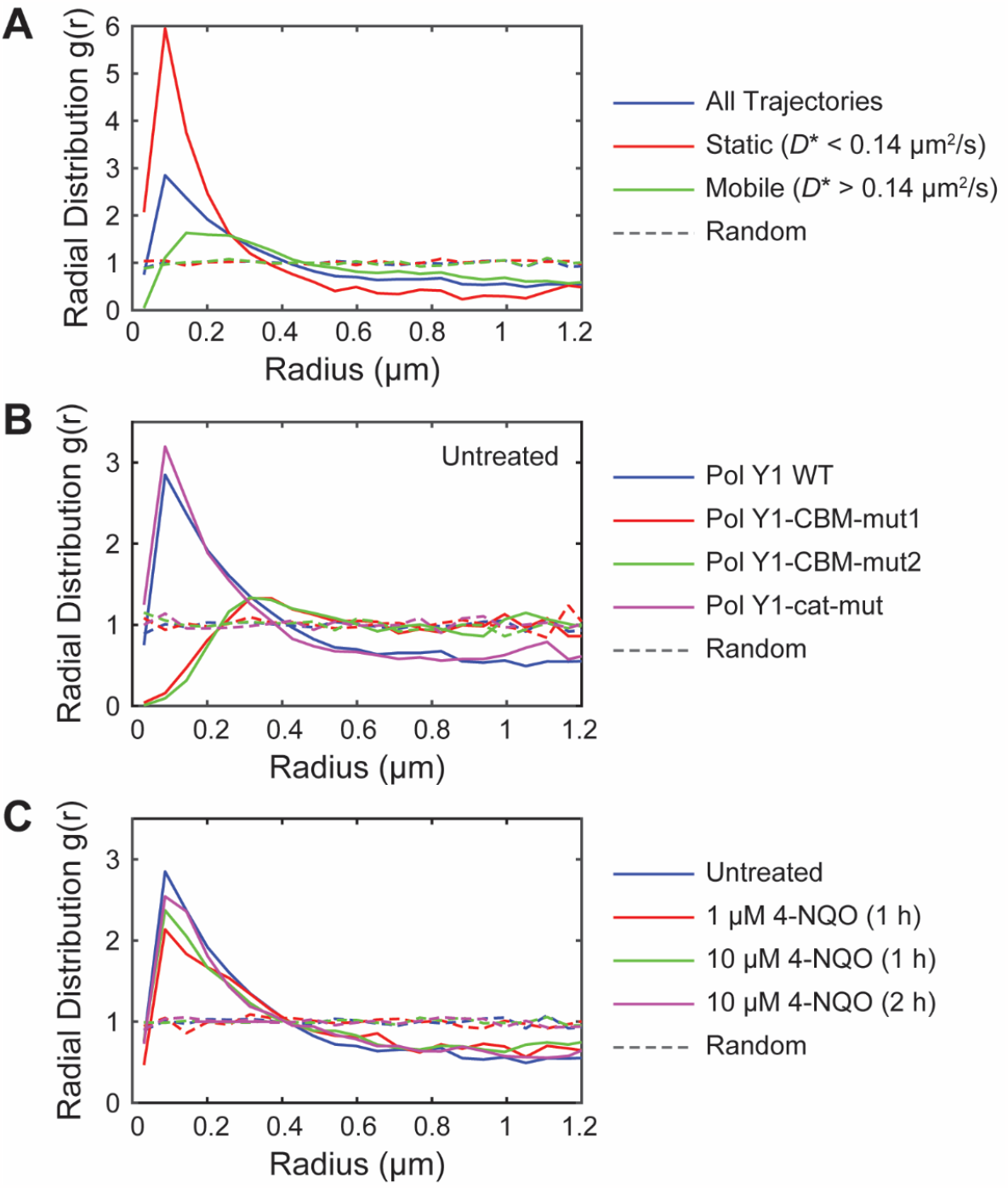
Radial distribution function *g*(*r*) analysis of Pol Y1-Halo and DnaX-mYPet colocalization. (A) Pol Y1-DnaX *g*(*r*) in untreated cells for all Pol Y1 trajectories, static Pol Y1 trajectories (*D*^*^ < 0.14 μm^2^/s), and mobile Pol Y1 trajectories (*D*^*^ > 0.14 μm^2^/s). (B) Pol Y1-DnaX *g*(*r*) for WT Pol Y1, Pol Y1-CBM-mut1, Pol Y2-CBM-mut2, and Pol Y1-cat-mut in untreated cells. (C) Pol Y1-DnaX *g*(*r*) in untreated cells, cells treated with 1 μM 4-NQO for 1 h, cells treated with 10 μM 4-NQO for 1 h, and cells treated with 10 μM 4-NQO for 2 h.

### There are static and mobile populations of Pol Y1 during normal growth

In *E. coli*, DNA polymerases exist in both static and mobile populations, characterized by slow and fast diffusion respectively.(14, 36) Next, we asked whether the same was true of Pol Y1 during normal growth. We quantified the motion of Pol Y1 by calculating an apparent diffusion coefficient, *D*^*^. The distribution of *D*^*^ values (Figure 5A) revealed a static population, with *D*^*^ ≈ 0, as well as a broad distribution of mobile molecules. To quantify these populations, we fit the *D*^*^ distribution to an analytical expression for three diffusing species. We found that approximately 28% of Pol Y1 molecules were static (*D*^*^ ≈ 0.08 μm^2^/s). In addition, the fit revealed two mobile populations. The more highly mobile population (*D*^*^ ≈ 0.98 μm^2^/s) represented 47% of the population, whereas the intermediate population (*D*^*^ ≈ 0.23 μm^2^/s) represented 25%. (See Table S5 for all diffusion coefficient distribution fit results.) The static population is immobilized through interactions with DNA, which diffuses very slowly in the cell, or with DNA-bound proteins; Pol Y1 molecules in this population may be, but are not necessarily, performing DNA synthesis. The more highly mobile population likely represents Pol Y1 molecules diffusing in the cytoplasm, whereas the intermediate population may represent transiently bound molecules.

**Figure 5.**
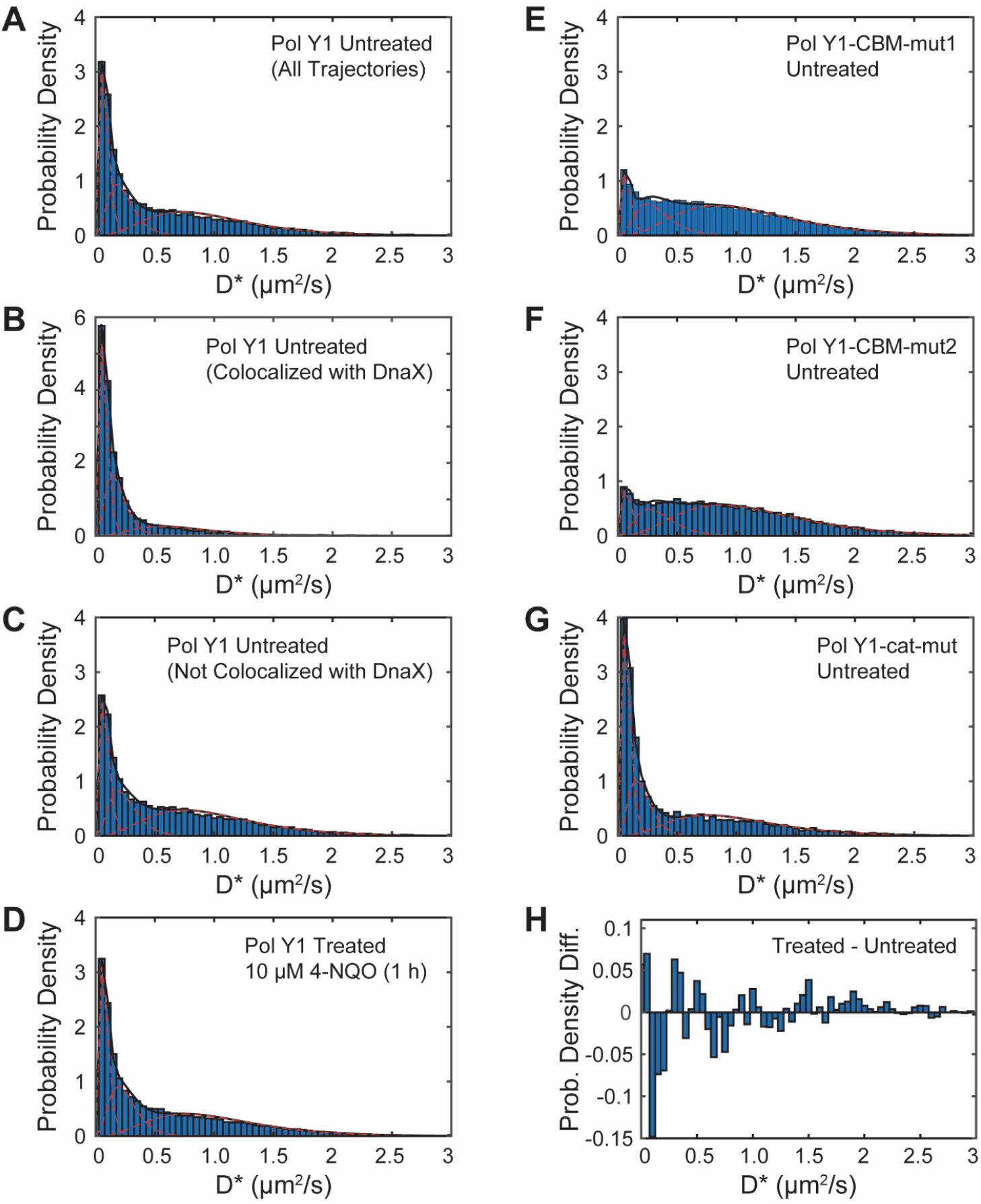
Apparent diffusion coefficient (*D*^*^) distributions for Pol Y1-Halo. *D*\* distributions for WT Pol Y1 in untreated cells for (A) all molecules, (B) molecules < 200 nm from DnaX-mYPet foci, and (C) molecules > 200 nm from DnaX-mYPet foci. (D) *D*\* distributions for WT Pol Y1 in cells treated with 10 μM 4-NQO for 1 h. *D*\* distributions for (E) Pol Y1-CBM-mut1, (F) Pol Y1-CBM-mut2, and (G) Pol Y1-cat-mut mutants in untreated cells. (H) Difference between *D*\* distributions for WT Pol Y1 in 4-NQO treated and untreated cells. Note different *y*-axis scales in panels (B) and (H).

Next, we asked whether the static Pol Y1 population was preferentially enriched near sites of replication. To address this question, we compared the *D*^*^ distributions for Pol Y1 molecules less than or greater than 200 nm from a DnaX focus. We found that Pol Y1 molecules localized near DnaX foci were more likely to be static (Figure 5B), whereas molecules at greater distances were slightly more likely to be mobile (Figure 5C), with the static population reflecting approximately 44% and 24% of the total, respectively. Consistent with this analysis, we found a greater degree of Pol Y1-DnaX colocalization (maximum *g*(*r*) ≈ 5.95) for static Pol Y1 molecules (*D*^*^ < 0.14 μm^2^/s) (Figure 4A). Mobile Pol Y1 molecules (*D*^*^ > 0.14 μm^2^/s) showed little to no colocalization at short *r* (*g*(*r*) ≈ 1.112 at the peak *r* position characteristic of all trajectories), reaching a slightly higher but still low level of colocalization (maximum *g*(*r*) ≈ 1.633) at slightly larger *r* values (Figure 4A). Taken together, these results are consistent with recruitment and possible activity of Pol Y1 at or near sites of replication during normal replication.

### The interaction with the DnaN clamp enriches Pol Y1 near sites of replication during normal growth

The activity of TLS polymerases in *E. coli* is mediated by critical protein-protein interactions with the β (DnaN) clamp; in particular, a strain bearing a Pol IV mutant unable to interact with the clamp is indistinguishable from a Pol IV knockout in damage tolerance.(14, 54) Pol Y1 contains a canonical CBM sequence (QLDLF) (Figure 1A),(18, 22) like the *E. coli* TLS polymerases, but little is known about the functional role of the clamp interaction. To investigate the role of clamp-binding in Pol Y1 activity, we introduced two different sets of mutations to the CBM. The first, designated CBM-mut1 (QADAF), was previously shown to reduce untargeted mutagenesis by Pol Y1.(18) The second, designated CBM-mut2 (ALDLA), contains mutations to the highly conserved Gln and Phe residues, which are predicted to weaken clamp-binding substantially.(22, 55) To test the functional consequence of these mutations, we assayed 4-NQO survival as described previously. Both mutants were as sensitized to 4-NQO treatment as the Pol IV knockout (Figure 2C), indicating that the clamp interaction is critical for Pol Y1 activity, like in *E. coli*.

Next, we investigated the effect of the clamp-binding mutations on the localization and dynamics of Pol Y1 during normal growth. In *E. coli*, the interaction with the clamp is essential for Pol IV activity,(56, 54, 14) but the interaction with SSB plays a greater role in Pol IV enrichment near sites of replication.(13, 15) We constructed C-terminal HaloTag fusions to both CBM mutants, in combination with the DnaX-mYPet replisome marker, and imaged cells under the same conditions as for WT Pol Y1. Radial distribution function analysis revealed a complete loss of Pol Y1-DnaX colocalization for both CBM mutants (*g*(*r*) ≈ 0.157 and 0.0938 at short *r* values for Pol Y1-CBM-mut1 and Pol Y1-CBM-mut2, respectively) (Figure 4B). In contrast to WT Pol Y1, the *g*(*r*) curves peaked at much larger *r* values, reaching (*g*(*r*) ≈ 1.3 at *r* > 300 nm; similar behavior has been observed for two proteins that are localized in similar regions of the cell without being truly colocalized.(14) This loss of colocalization with DnaX for the Pol Y1 CBM mutants is consistent with a reduction in their average localization at the long axis quarter and three-quarter positions (Figures S3A and C). Likewise, there was a substantial reduction in the static population of Pol Y1 (*D*^*^ ≈ 0.08 μm^2^/s), from 28% for the Pol Y1 WT to 11% and 9% for Pol Y1-CBM-mut1 (Figure 5E) and Pol Y1-CBM-mut2 (Figure 5F), respectively. Thus, the clamp interaction appears to play a major role in immobilizing and enriching Pol Y1 near sites of replication in *B. subtilis*.

### Pol Y1 catalytic activity is not required for enrichment near sites of replication

Like other DNA polymerases, Pol Y1 contains conserved amino acids with negatively charged side chains that coordinate catalytic Mg^2+^ ions during DNA synthesis.(57) To investigate the effect of Pol Y1 catalytic activity on its localization and dynamics in vivo, we introduced the catalytically inactivating D108 E109 → A108 A109 mutations.(18) Pol Y1 catalytic activity was previously shown to be required for untargeted mutagenesis,(18) but the effect on DNA damage tolerance has not been explored. First, we tested the survival of the Pol Y1-cat-mut catalytically inactive mutant in response to 4-NQO treatment. Like the clamp-binding mutants, the catalytic mutant was as sensitized to 4-NQO as the Pol Y1 knockout, indicating that catalytic activity is essential for damage tolerance (Figure 2C).

To look for any effects of catalytic activity on Pol Y1 localization and dynamics, we imaged cells with a Pol Y1-Halo fusion bearing the catalytic mutation during normal growth. In contrast to the results for the CBM mutants, radial distribution function analysis revealed a small increase in Pol Y1-DnaX enrichment for the catalytically inactive mutant relative to WT (maximum *g*(*r*) ≈ 3.20) (Figure 4B), consistent with a similar average cellular localization pattern (Figure S3E). There was likewise a slight increase in the static population of Pol Y1 molecules (*D*^*^ ≈ 0.08 µm^2^/s), from 28% for the Pol Y1 WT to 34% for Pol Y1-cat-mut (Figure 5G). These results demonstrate that catalytic activity is not required to stabilize Pol Y1 near sites of replication. Instead, the modest increases in colocalization with sites of replication and in the static population is consistent with a picture in which Pol Y1-cat-mut spends more time bound near sites of replication due to its inability to perform DNA synthesis.

To compare the dynamics of the catalytically inactive mutant and WT Pol Y1 more carefully, we measured the binding lifetime for static Pol Y1 molecules imaged using a long 250 ms integration time. In this approach, the apparent binding lifetime is taken as the duration of the trajectory; when the Pol Y1 molecule dissociates from DNA or from a DNA-bound protein, the trajectory ends (Figure S6A). However, photobleaching of the JFX_554_ fluorophore also terminates the trajectory and cannot be distinguished from a true dissociation event (Figure S6A). To measure the JFX_554_ photobleaching rate under our imaging conditions, we imaged JFX_554_-labeled WT Pol Y1 in fixed cells, where Pol Y1 is immobilized and there is no contribution from dissociation.(14, 36) Comparison of the apparent binding lifetime for WT Pol Y1 in untreated and fixed cells (Figure S6B) revealed similar short-timescale dynamics, but a longer binding lifetime in fixed cells at longer timescales. In contrast, Pol Y1-cat-mut showed an intermediate binding lifetime at longer timescales, consistent with the small increase in the static population observed in diffusion coefficient measurements. To obtain photobleaching-corrected binding lifetime estimates, we first fit the apparent binding lifetime distributions to exponential functions (Figures S6D – F and Table S6). From the fit parameters, we extracted an effective photobleaching lifetime (1.10 ± 0.05 s) and photobleaching-corrected binding lifetimes for WT Pol Y1 (1.9 ± 0.4 s) and Pol Y1-cat-mut (6 ± 3 s), showing an approximately 3-fold increase in the binding lifetime for the catalytically inactive mutant relative to WT.

### DNA damage leads to minimal changes in Pol Y1 localization and dynamics

In *E. coli*, the Pol Y1 homolog Pol IV is recruited to sites of replication in response to a variety of replication perturbations, including treatment with cognate DNA damaging agents, which generate DNA lesions that Pol IV can bypass efficiently.(14, 15) Given that Pol Y1 colocalizes with sites of replication during normal growth, we next asked whether this enrichment would increase upon DNA damage. To address this question, we chose to investigate the response of Pol Y1 to treatment with the drug 4-NQO; as shown previously, Pol Y1 promotes cell survival upon 4-NQO treatment.

First, we determined a 4-NQO treatment condition that would have a moderate effect on cell viability without being highly lethal. We tested exposure to a 1 μM 4-NQO concentration for 1 h in liquid culture and quantified the fold-change in the number of CFUs/mL by plating cells before and after treatment and counting colonies (Table S7). The number of CFUs/mL increased by a factor of 2.1 ± 0.2 (mean ± std.) after this treatment. In comparison, the number of CFUs/mL increased by 2.3 ± 0.6 in cells treated with the solvent DMF alone and by 2.09 ± 0.08 in untreated cells, indicating that this 4-NQO treatment had little or no effect on cell growth. We next tested higher concentrations of 10 and 50 μM with the same 1 h incubation time. For 10 μM, we observed a slight decrease in the number of CFUs/mL (fold change of 0.7 ± 0.2), whereas the 50 µM concentration produced a substantial decrease (fold change of 0.06 ± 0.02). Although these 4-NQO concentrations are one to two orders of magnitude greater than the concentrations used in the survival assays on solid media, it should be noted that the duration of treatment is much shorter (1 h in comparison to approximately 16 h).

In light of these results, we chose the 10 μM 4-NQO concentration as a starting point. We imaged cells after 1 h of treatment and quantified the localization and dynamics of Pol Y1. First, we looked at changes in the number and localization of DnaX foci. There was a small increase in the number of DnaX foci per cell (mean ± S.E.M.: 2.00 ± 0.04, *p* < 10^−5^) in comparison to untreated cells. Although foci were still primarily localized between the quarter and three-quarter positions along the long cell axis, there was increased localization at midcell (Figures 3D and S2D). Similar behavior has been observed in *E. coli* and may reflect a slowdown in replication or a failure to initiate subsequent rounds of replication.(14) Notably, we observed analogous changes in Pol Y1 localization, with an increase in the fraction of Pol Y1 molecules at the midcell position along the long cell axis (Figures 3E and S2F). Consistent with these coupled changes in both DnaX and Pol Y1 localization, 4-NQO treatment had little effect on Pol Y1-DnaX colocalization. The radial distribution function showed moderate colocalization, with a slight reduction (maximum *g*(*r*) ≈ 2.370) relative to untreated cells (Figure 4C).

In addition to the minimal changes in Pol Y1-DnaX localization, we also observed little change in Pol Y1 mobility upon 4-NQO treatment. The distribution of *D*^*^ values (Figure 5D) was comparable to that in untreated cells (Figure 5A), with approximately 28% of Pol Y1 molecules in the static (*D*^*^ ≈ 0.08 μm^2^/s) population; the change in the *D*^*^ distribution upon treatment is shown in Figure 5H. Likewise, we observed only a minimal change in the Pol Y1 binding lifetime upon 4-NQO treatment, with similar dynamics at both short and long timescales (Figure S6C). Fitting the apparent binding lifetime distribution (Figure S6G) yielded a photobleaching corrected binding lifetime of 2.2 ± 0.5 s, similar to the lifetime of 1.9 ± 0.4 s measured in untreated cells. Taken together, these results indicate that the dynamics of Pol Y1 do not change substantially under this 4-NQO treatment condition.

To explore a possible dose-dependence in the response of Pol Y1 to 4-NQO DNA damage, we tested treatment with a lower 1 μM concentration for 1 h. As expected, we saw little effect on Pol Y1-DnaX colocalization (Figure 4C) or Pol Y1 mobility (Figure S5E) for treatment with this lower concentration. Because cell viability was substantially reduced at higher 4-NQO concentrations, we did not test increased doses. Instead, we looked for possible delayed responses to treatment by exposing cells to 10 μM 4-NQO for a longer 2 h incubation, which slows cell growth but does not lead to a loss of viability (fold change of 1.5 ± 0.3 in the number of CFUs/mL). Again, we observed minimal changes in Pol Y1-DnaX colocalization (Figure 4C) or in the Pol Y1 diffusion coefficient (Figure S5F) upon this treatment. In both cases, however, the average Pol Y1 localization pattern was closer to that in untreated cells (Figures S2G and H). Thus, we were unable to find a 4-NQO treatment condition that led to substantial recruitment of Pol Y1 to sites of replication, as observed for *E. coli* Pol IV, or to substantial changes in Pol Y1 dynamics.

In addition to characterizing the response of WT Pol Y1 to 4-NQO treatment, we also determined the response of the clamp-binding and catalytically inactive mutants. Overall, the mutant Pol Y1 localization (Figures S3B, D, and F), colocalization with sites of replication (Figure S4C), and diffusion (Figures SG – I) were consistent with the observed behavior for each mutant in untreated cells. The catalytically inactive mutant, Pol Y1-cat-mut, showed modest increases in colocalization with sites of replication (Figure S4C) and in the fraction of static molecules (Figure S5I) relative to WT Pol Y1. In contrast, the clamp-binding mutants, Pol Y1-CBM-mut1 and Pol Y1-CBM-mut2, were not colocalized with sites of replication (Figure S4C) and had substantial reductions in the static Pol Y1 population (Figures S5G and H). Thus, 4-NQO treatment does not appear to alter the molecular requirements for Pol Y1 enrichment and stabilization at sites of replication; the clamp-binding interaction is still required, but catalytic activity is not.

## Discussion

Y-family DNA polymerases and the TLS pathway are conserved across a wide range of bacterial species, yet most previous work has focused on the model gram-negative bacterium *E. coli*.(3) In this study, we used single-molecule fluorescence microscopy to characterize the localization and dynamics of the TLS polymerase Pol Y1, a homolog of *E. coli* Pol IV, in the gram-positive bacterium *B. subtilis*. We created a new functional HaloTag fusion to the endogenous copy of Pol Y1 and used quantitative fluorescence microscopy and single-particle tracking to determine its dynamics and localization at the single-molecule level in live cells, both in the absence of exogenous DNA damage and upon treatment with the drug 4-NQO.

Given the low fidelity and slow rate of synthesis of most TLS polymerases, their access to the DNA template during normal replication can be harmful.(58) Although TLS polymerases promote mutagenesis if overexpressed,(18, 59) in *E. coli* they make little contribution to the basal mutation rate at normal expression levels.(60) An emerging model suggests an explanation for these observations in terms of a spatial component to the regulation of TLS polymerases in *E. coli*; in this model, TLS polymerases are restricted from the replication fork during normal growth and only enriched, if at all, upon perturbations to replication (Figure 6, top panels).(10, 13–16) In contrast, we found that *B. subtilis* Pol Y1 is moderately enriched at or near sites of replication during normal growth (Figure 6, bottom left panel). By measuring the diffusion of Pol Y1 molecules, we showed that the static Pol Y1 population is particularly enriched at these sites, suggesting that Pol Y1 is interacting either with DNA or with other DNA-bound proteins at or near the replication fork. Unlike *E. coli* TLS polymerases, Pol Y1 is not upregulated by the SOS DNA damage response.(18, 21) Taken together with our imaging results, this observation supports the possibility that Pol Y1 is playing a role in normal replication, possibly in alleviating replication-transcription conflicts or in the transcription-coupled NER pathway.(25) Notably, the transcription/repair coupling factor Mfd is found to be associated with the nucleoid even in the absence of exogenous DNA damage in *E. coli*.(61) More broadly, these results indicate that there are different patterns of spatial regulation of TLS polymerases across different bacterial species and suggest that studies in other species may reveal different mechanisms.

**Figure 6.**
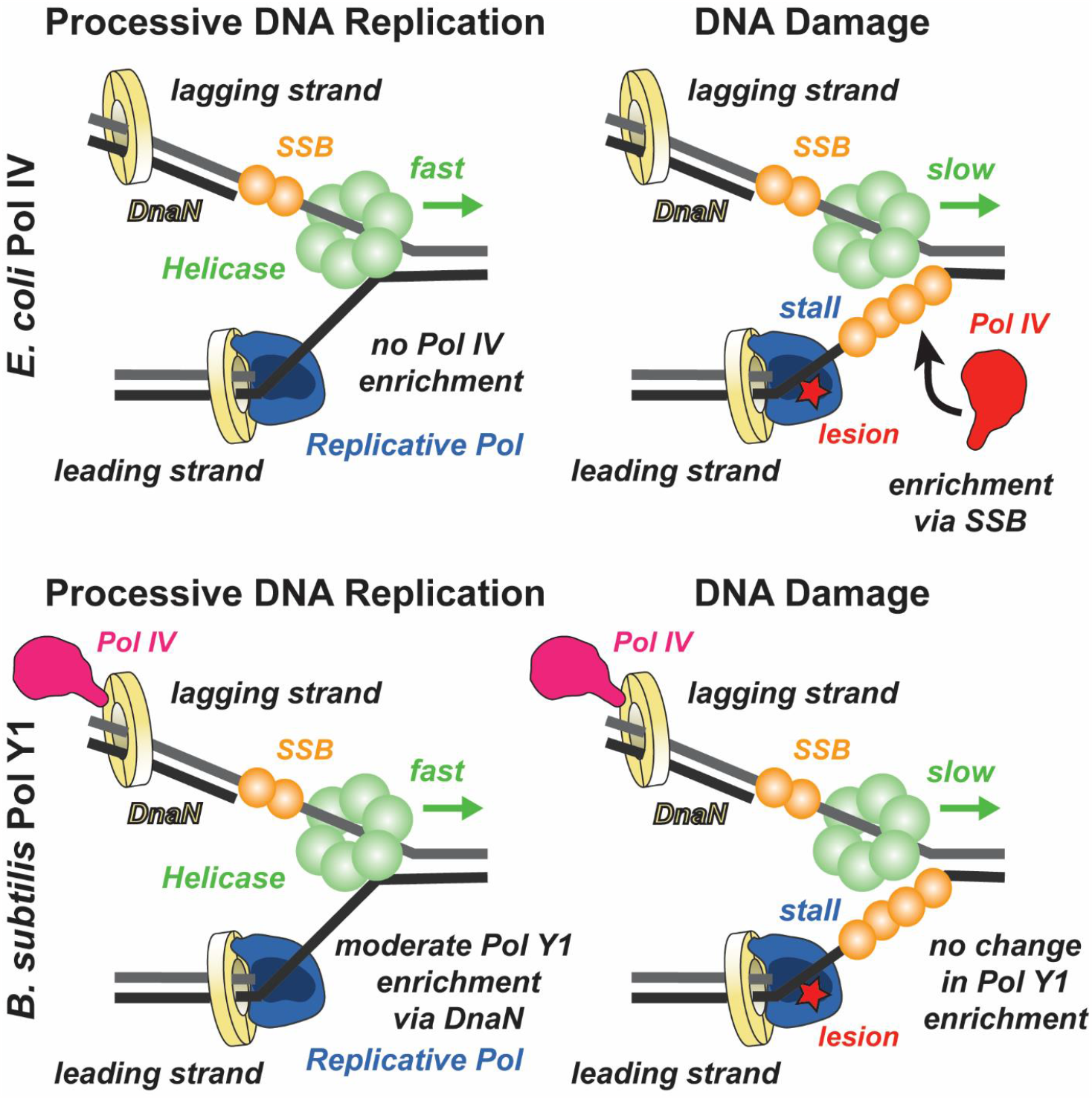
Model of different TLS polymerase recruitment mechanisms in *E. coli* and *B. subtilis*. In *E. coli*, there is little or no enrichment of Pol IV near the replication fork during processive replication (*top left*), but a significant increase in enrichment upon DNA damage-induced replication stalling (*top right*), primarily through interactions with SSB. In *B. subtilis*, Pol Y1 is moderately enriched near the replication fork through interactions with DnaN, both during processive replication (*bottom left)* and upon DNA damage-induced replication stalling (*bottom right*).

In *E. coli*, changes in the localization and dynamics of TLS polymerases have been observed upon DNA damage or other perturbations to replication; in particular, Pol IV is strongly recruited to sites of replication upon induction of DNA damage by methyl methanesulfonate (MMS) or ultraviolet (UV) light (Figure 6, top right panel).(15) In this study, we characterized the localization and dynamics of Pol Y1 after different treatments with the drug 4-NQO; consistent with previous work,(25) we found that Pol Y1, but not Pol Y2, mediates cell survival to 4-NQO treatment. Surprisingly, we observed little or no change in Pol Y1 behavior, including its colocalization with sites of replication and its dynamics, upon 4-NQO treatment (Figure 6, bottom right panel). In contrast to Pol IV, Pol Y1 is not transcriptionally upregulated by the SOS DNA damage response.(18, 21) Although SOS induction of Pol IV plays an important role in its response to DNA damage, selective recruitment of Pol IV to the replication fork upon replication stalling was observed in a constitutively SOS-induced strain where Pol IV levels were unchanged upon DNA damage.(14, 15) Thus, the lack of SOS induction of Pol Y1 is likely not sufficient to explain the lack of response to 4-NQO treatment. These results raise the possibility that Pol Y1 promotes cell survival upon DNA damage through different mechanisms than Pol IV.

Like replicative DNA polymerases, TLS polymerases function with the assistance of accessory proteins. In *E. coli*, the interaction between TLS polymerases and the replication processivity factor, the sliding clamp, is essential for TLS.(3, 54, 56) Recent single-molecule imaging experiments have revealed that the clamp interaction contributes to the enrichment of Pol IV at sites of replication upon DNA damage.(14) However, interactions with another replication protein, SSB, are the primary driver for Pol IV localization (Figure 6, top right panel).(13) Like Pol IV, Pol Y1 contains a conserved CBM, suggesting that it interacts with the DnaN clamp.(18) We characterized two Pol Y1 mutants with modifications to its CBM that are predicted to eliminate clamp binding and found that both mutants were as sensitive to treatment with 4-NQO as was a Pol Y1 knockout, indicating that the clamp interaction is critical for Pol Y1-mediated damage tolerance. Single-molecule imaging of these CBM mutants revealed a reduction in the static population of Pol Y1 and a loss of Pol Y1-DnaX colocalization, both during normal growth and upon 4-NQO treatment. Taken together, these results indicate that Pol Y1 is interacting with DNA-bound clamps at or near the replication fork (Figure 6, bottom panels), but they do not exclude the possibility that other protein-protein interactions, particularly with SSB, play a role in Pol Y1 activity.

We also investigated the effect of Pol Y1 catalytic activity on its localization and dynamics. In *E. coli*, imaging studies of Pol IV have reached different conclusions about the role of catalytic activity in localization; one found a complete loss of Pol IV foci upon DNA damage for a catalytically inactive mutant,(10) whereas another found only a modest decrease in Pol IV colocalization with sites of replication.(14) Here, we observed that a catalytically inactive Pol Y1 mutant, although broadly similar to WT Pol Y1 in its behavior, showed several hallmarks of increased stability at sites of replication. Although the changes were modest, Pol Y1-cat-mut had greater enrichment at sites of replication relative to WT Pol Y1 and a larger static population in diffusion coefficient measurements, both during normal growth and upon 4-NQO treatment. Likewise, there was an increase in the binding lifetime of static Pol Y1-cat-mut molecules relative to WT during normal growth. These results support a picture where stabilization of Pol Y1 near the replication fork does not require DNA synthesis.

Taken together, our results show that there are significant differences in the activity of *B. subtilis* Pol Y1 in comparison to its *E. coli* homolog Pol IV (Figure 6). This work is the first step toward elucidating the mechanisms of TLS polymerases in *B. subtilis* at the single-molecule level, but many key questions remain. First, is the lack of damage-induced Pol Y1 enrichment at sites of replication unique to 4-NQO treatment, or is it common to all types of DNA damage? Although beyond the scope of this work, future studies should explore the response of Pol Y1 to different types of replication perturbations, including diverse forms of DNA damage. Second, what other protein-protein interactions, if any, play a role in Pol Y1 activity? We have shown that the interaction with the DnaN clamp is essential to 4-NQO damage tolerance and drives Pol Y1 localization and dynamics in vivo. Although at least 12 SSB-interacting proteins have been identified in *B. subtilis*, Pol Y1 is not among them.(19, 62) By analogy to the *E. coli* TLS polymerases, however, the possibility of an interaction with SSB should be explored. Additionally, there is experimental or computational evidence for interactions with RecA(24) or Pol I(23) that merit further investigation. Finally, what are the primary cellular functions of Pol Y1? Its lack of SOS induction and constitutive enrichment near sites of replication suggest that it may have an important role beyond replication-coupled TLS. In particular, future studies should explore Pol Y1 involvement in resolving replication-transcription conflicts, including a possible role in the transcription-coupled NER pathway. Answering these and other questions about TLS polymerase activity in *B. subtilis* will help broaden bacterial models for TLS beyond *E. coli* and provide new insight into how cells maintain a balance between genome stability and mutagenesis under stress.

## Data Availability

Data and custom MATLAB code from this study are available in the Zenodo repository (DOI: 10.5281/zenodo.8411186).

## Acknowledgments

We thank Xindan Wang (Indiana University) for sharing several bacterial strains and for advice in strain construction, Joseph Loparo (Harvard Medical School) for sharing bacterial strains, and Luke Lavis (Howard Hughes Medical Institute Janelia Research Campus) for providing Janelia Fluor dyes. We also acknowledge Nicholas Bafundo for contributions to the strain construction strategy and Emily Holmes and Luke O’Neal for assistance with microscopy.

