## Supplementary Information for "“The Translesion Polymerase Pol Y1 is a Constitutive Component of the *B. subtilis* Replication Machinery”"

**This PDF file includes:**

Figures S1 to S6
Tables S1 to S7
Supplementary Methods
Supplementary References

### Supplementary Figures

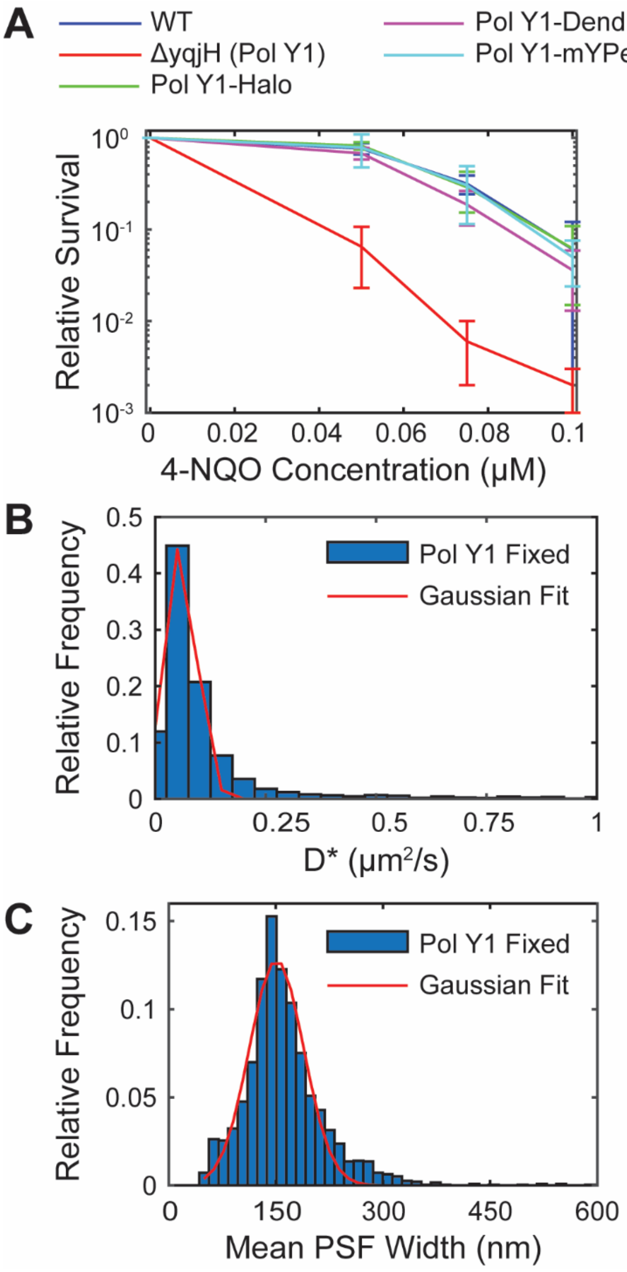


**Figure S1.** (A) Relative survival of *B. subtilis* strains treated with different concentrations of 4-NQO: WT Pol Y1, Pol Y1 knockout, Pol Y1-Halo fusion, Pol Y1-Dendra2 fusion, and Pol Y1-mYPet fusion strains. (B) Distribution of the Pol Y1 apparent diffusion coefficient $D^*$ measured in fixed cells and the corresponding Gaussian fit. (C) Distribution of the mean point spread function (PSF) width for static trajectories recorded with a long 250 ms integration time and the corresponding Gaussian fit.

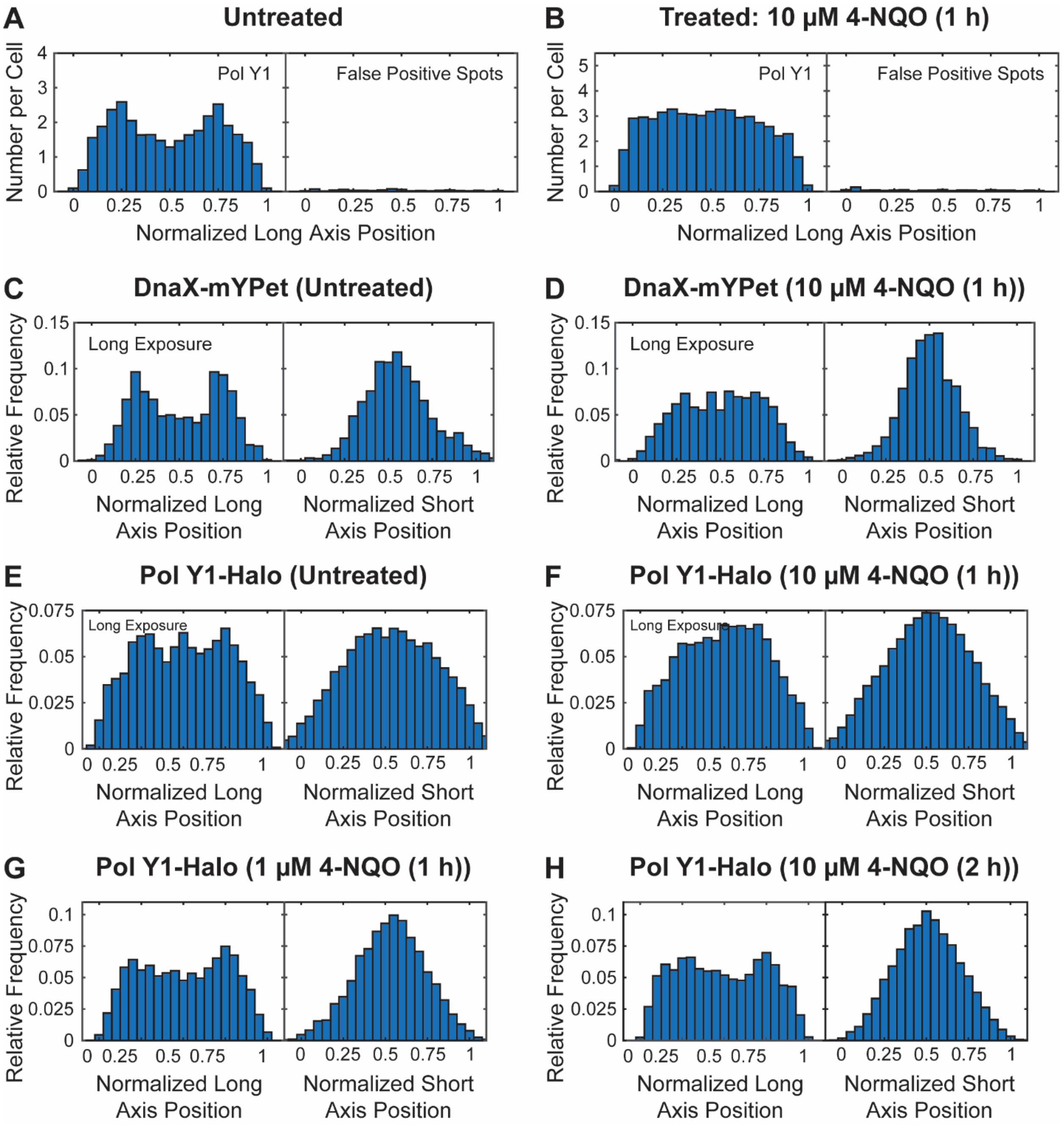


**Figure S2.** Cellular localization of DnaX-mYPet and Pol Y1-Halo. Long cell axis projections of Pol Y1 (left) and false positive spots (right) in (A) untreated cells and (B) cells treated with 10 μM 4-NQO for 1 h on a per cell basis. Long and short cell axis projections of (C, D) DnaX and (E, F) Pol Y1 in untreated cells and cells treated with 10 μM 4-NQO for 1 h, respectively, recorded with a long 250 ms integration time. Long and short cell axis projections of Pol Y1 in cells treated with (G) 1 μM 4-NQO for 1 h and (H) 10 μM 4-NQO for 2 h.

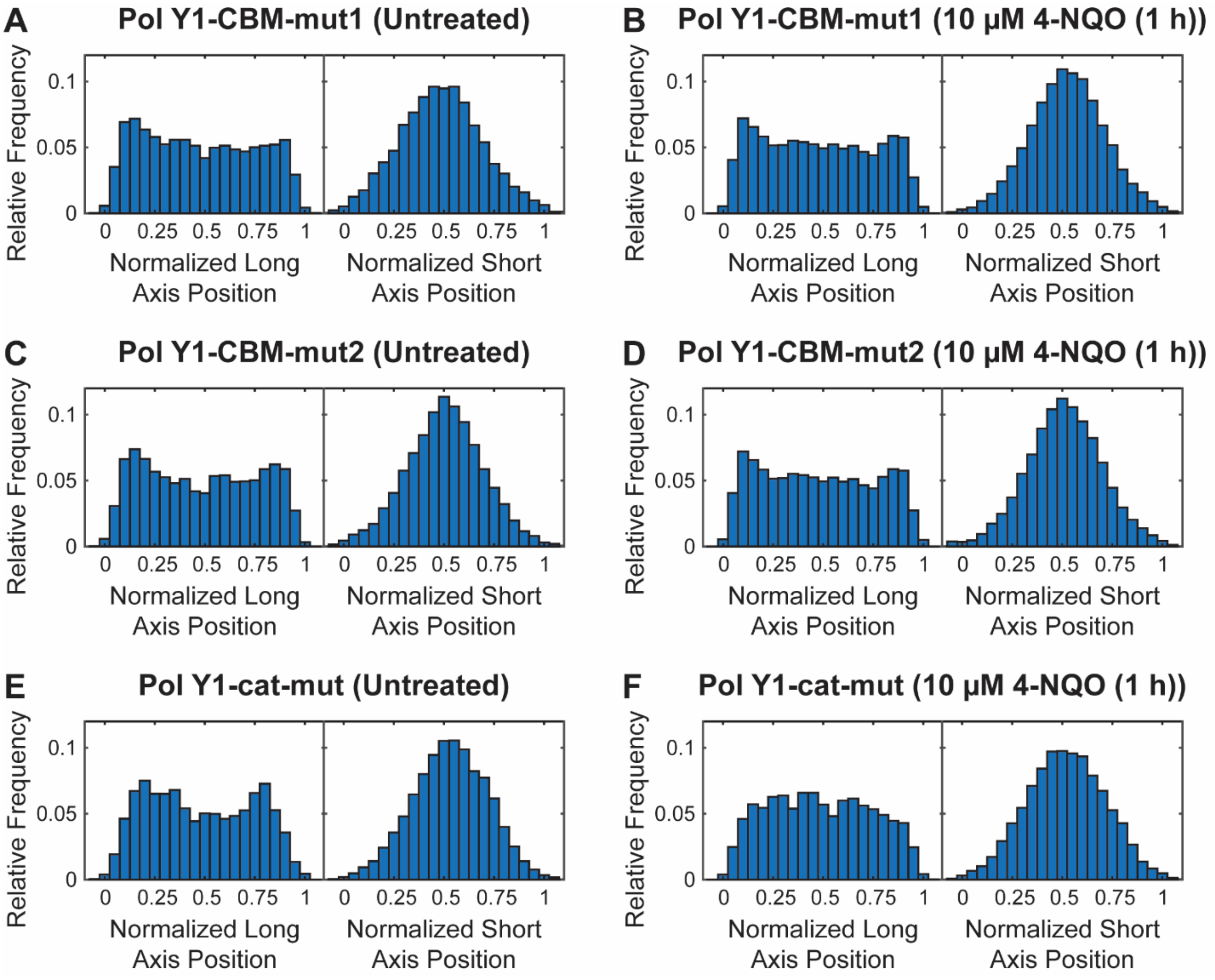


**Figure S3.** Cellular localization of Pol Y1-Halo mutants. Long and short cell axis projections of Pol Y1-CBM-mut1, Pol Y1-CBM-mut2, and Pol Y1-cat-mut in (A, C, E) untreated cells and (B, D, F) cells treated with 10 μM 4-NQO for 1 h, respectively.

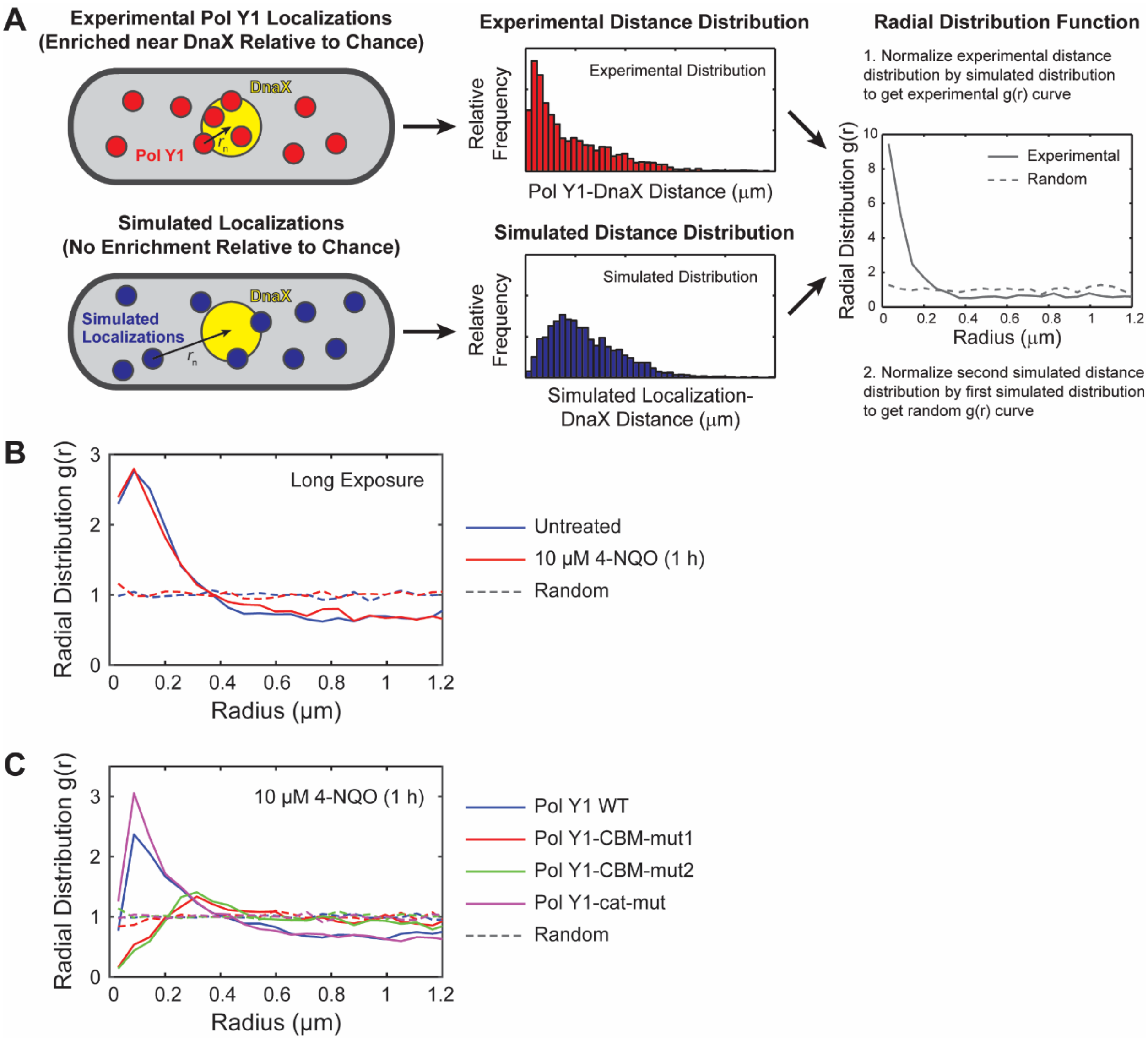


**Figure S4.** Radial distribution function $g(r)$ analysis of Pol Y1-Halo and DnaX-mYPet colocalization. (A) Cartoon of radial distribution function $g(r)$ analysis. (B) Pol Y1-DnaX $g(r)$ in untreated cells and cells treated with 10 μM 4-NQO for 1 h recorded with a long 250 ms integration time. (C) Pol Y1-DnaX $g(r)$ for WT Pol Y1, Pol Y1-CBM-mut1, Pol Y2-CBM-mut2, and Pol Y1-cat-mut in cells treated with 10 μM 4-NQO for 1 h.

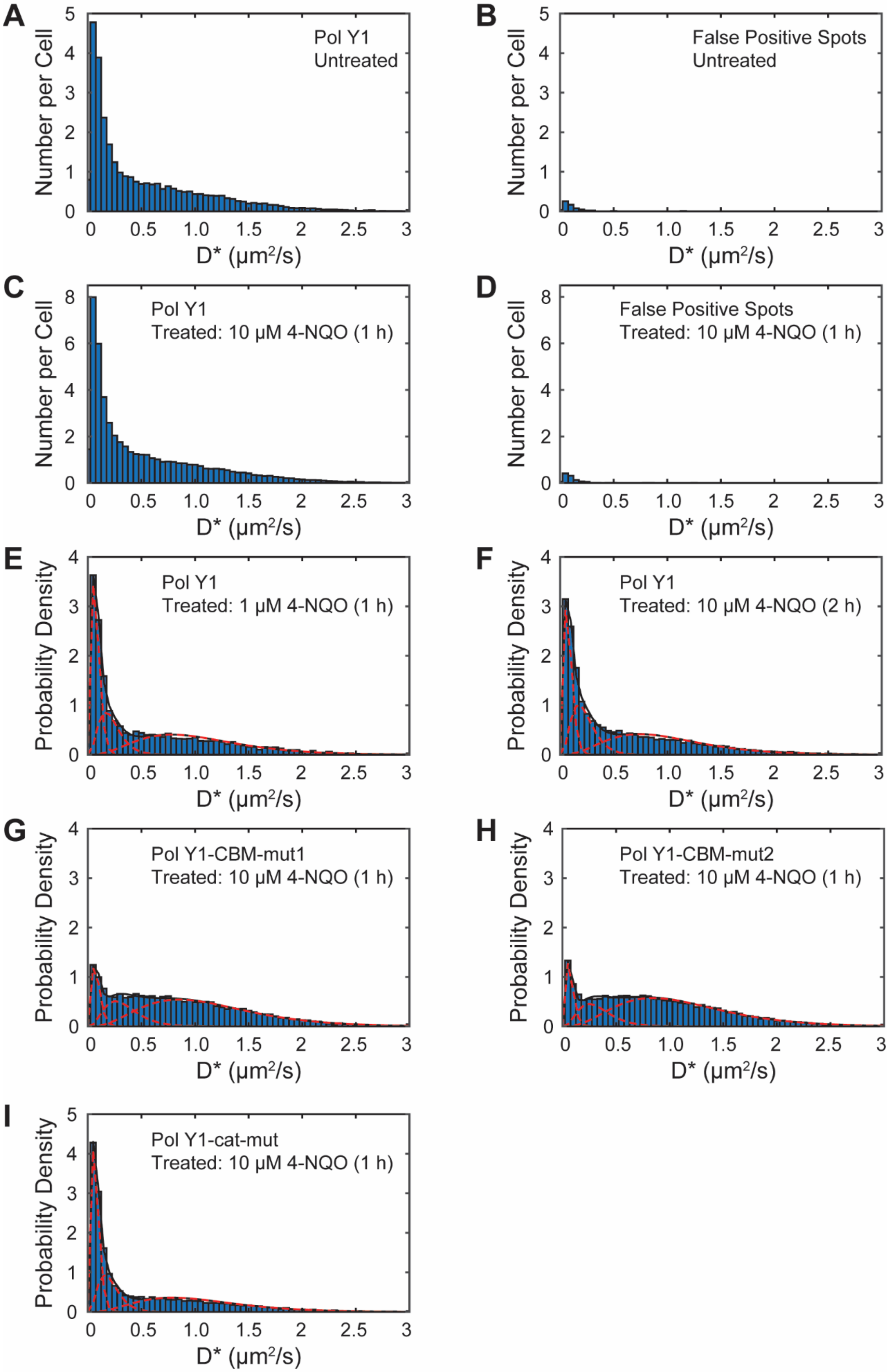
A
Number per Cell
Pol Y1
Untreated
D* (μm²/s)
B
False Positive Spots
Untreated
C
Pol Y1
Treated: 10 μM 4-NQO (1 h)
D
False Positive Spots
Treated: 10 μM 4-NQO (1 h)
E
Probability Density
Pol Y1
Treated: 1 μM 4-NQO (1 h)
F
Pol Y1
Treated: 10 μM 4-NQO (2 h)
G
Pol Y1-CBM-mut1
Treated: 10 μM 4-NQO (1 h)
H
Pol Y1-CBM-mut2
Treated: 10 μM 4-NQO (1 h)
I
Pol Y1-cat-mut
Treated: 10 μM 4-NQO (1 h)

**Figure S5.** Apparent diffusion coefficient ($D^*$) distributions for Pol Y1-Halo. $D^*$ distributions for Pol Y1 (A, C) and false positive spots (B, D) in untreated cells and cells treated with 10 μM 4-NQO for 1 h, respectively, on a per cell basis. $D^*$ distributions for WT Pol Y1 in cells treated with (E) 1 μM 4-NQO for 1 h and (F) 10 μM 4-NQO for 2 h. $D$* distributions for (G) Pol Y1-CBM-mut1, (H) Pol Y1-CBM-mut2, and (I) Pol Y1-cat-mut mutants in cells treated with 10 μM 4-NQO for 1 h.

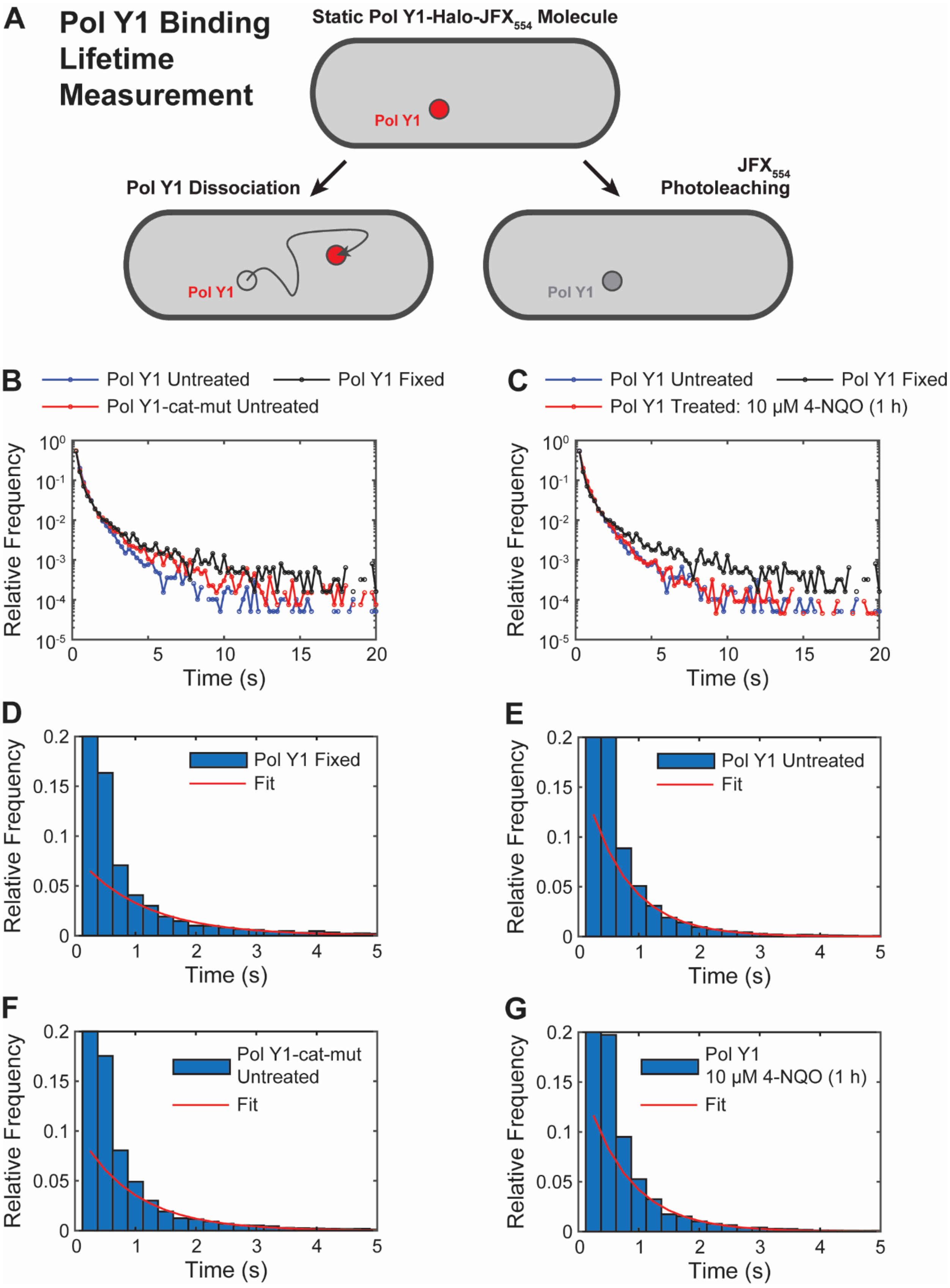
A
Pol Y1 Binding Lifetime Measurement
Static Pol Y1-Halo-JFX554 Molecule
Pol Y1
Pol Y1 Dissociation
JFX554 Photoleaching
B
Pol Y1 Untreated
Pol Y1 Fixed
Pol Y1-cat-mut Untreated
Relative Frequency
Time (s)
C
Pol Y1 Treated: 10 µM 4-NQO (1 h)
D
E
F
G
Fit
Pol Y1
10 µM 4-NQO (1 h)

**Figure S6.** Pol Y1 binding lifetime measurements. (A) Cartoon of Pol Y1 dissociation and $JFX_{554}$ photobleaching pathways. (B) Apparent Pol Y1 binding lifetime for WT Pol Y1 in untreated cells, Pol Y1-cat-mut in untreated cells, and WT Pol Y1 in cells fixed with formaldehyde. (C) Apparent Pol Y1 binding lifetime for WT Pol Y1 in untreated cells, cells treated with 10 μM 4-NQO for 1 h, and cells fixed with formaldehyde. Distributions of apparent binding lifetime and corresponding exponential fits for (D) WT Pol Y1 in fixed cells, (E) WT Pol Y1 in untreated cells, (F) Pol Y1-cat-mut in untreated cells, and (G) WT Pol Y1 in cells treated with 10 μM 4-NQO for 1 h. (Note that the *y*-axes are truncated in D – G to show the longer timescale behavior more clearly.)

### Supplementary Tables

### Table S1: Oligonucleotides used in this study

| Number | Designation | Sequence (5′-3′) |
|---|---|---|
| oEST008 | loxP ab rev univ short | GACCAGGGAGCACTGGTCAAC |
| oEST023 | yqjH-amplify-for | GCGGGTTGATATTATGCTCCTG |
| oEST028 | yqjH-downstream-rev | CGACAACTTCAGATGGGCTGGTGTTC |
| oEST029 | yqjH-amplify-rev-nostop | GCTTTTCTTTTCATCTTGAA |
| oEST030 | yqjH-linker-halo-for | ttcaagatgaaaagaaaagcGGCTCTGGACAGGGCTCAGG |
| oEST031 | halo-spec-rev | gcgagggagcagaaggatccTTACTAGCCGCTGATTTCTA |
| oEST032 | spec-for | GGATCCTTCTGCTCCCTCGCT |
| oEST033 | yqjH-linker-dendra2-for | ttcaagatgaaaagaaaagcCTCGAGGGATCTGGCGGATC |
| oEST034 | dendra2-spec-rev | gcgagggagcagaaggatccCTACCAGACTTGTGACGGCA |
| oEST035 | yqjH-linker-mYPet-for | ttcaagatgaaaagaaaagcCTCGAGGGATCAGGACAGGG |
| oEST036 | mYPet-spec-rev | gcgagggagcagaaggatccTTACTTGTAAAGTTCATTCA |
| oEST037 | yqjH-upstream-for | CATCAGTCACCGTATTGACT |
| oEST038 | yqjH-Nter-rev | TCGGCTCTTTCCCGGCATAA |
| oEST039 | yqjH-Nter-spec-iso-for | ttatgccgggaaagagccgaGGATCCTTCTGCTCCCTCGC |
| oEST040 | yqjH-Cter-spec-iso-rev | attcagcttttcttttcatcGACCAGGGAGCACTGGTCAA |
| oEST041 | yqjH-Cter-for | GATGAAAAGAAAAGCTGAATCGC |
| oEST048 | yqjW-downstream-rev | GTTCATCAAATTGGCTCACG |
| oEST049 | yqjW-upstream-for | AATATAAATCGGCCGGCCAG |
| oEST058 | yqjH-CBM-mut1-rev | TGAACgcATCGgcCTGTTTATAGGCCTGCTCTTTTTCTAC TAAATCCG |
| oEST059 | yqjH-CBM-mut1-for | CCTATAAACAGgcCGATgcGTTCAGCTTTAATGAAGATGC GAAGGAT |
| oEST060 | yqjH-CBM-mut2-rev | TGgcCAAATCGAGCgcTTTATAGGCCTGCTCTTTTTCTAC TAAATCCG |
| oEST061 | yqjH-CBM-mut2-for | CCTATAAAgcGCTCGATTTGgcCAGCTTTAATGAAGATGC GAAGGAT |
| oEST062 | yqjH-cat-mut-rev | CCATATAGCCTgCGgCGATGGAGACAGGCTCCACTAGGTC AGTATATT |
| oEST063 | yqjH-cat-mut-for | TCTCCATCGcCGcAGGCTATATGGACATGACCGATACA |
| oEST070 | yqjH-amplify-rev-iso | ctgagcgagggagcagaaggatccTCAGCTTTTCTTTTCA TCTTGAAA |
| oEST071 | yqjH-downstream-for-iso | gtagttgaccagtgctccctggtcATCGCTTGAAAAAAAG GGTG |
| oEST072 | yqjW-Nter-rev-iso | ctgagcgagggagcagaaggatccCACTTTTTCTTTCATC ATACACACC |
| oEST073 | yqjW-Cter-for-iso | gtagttgaccagtgctccctggtcAAAATAGGGGGGCATT ATAAA |

**Table S2: *B. subtilis* bacterial strains used in this study**

| Number | Designation or description | Relevant genotype | Construction or source strain designation | Reference |
|---|---|---|---|---|
| EST003 | *B. subtilis* prototrophic wild-type strain | PY79 | — | (1, 2) |
| EST053 | DnaX-mYPet | PY79 *dnaX-mYpet cat Ω pWX340a* | Gift of Xindan Wang (Indiana University); strain BWX519 from plasmid pWX340 | (3) |
| EST057 | PolC-mYPet | PY79 *polC-mYpet cat* | Gift of Xindan Wang (Indiana University); strain BWX499 | — |
| EST081 | PolC-Dendra2 | PY79 *polC-dendra2 loxP spec* | Gift of Xindan Wang (Indiana University); strain BWX2913 | — |
| EST083 | PolC-HaloTag | PY79 *polC-halo loxP spec* | Gift of Joseph Loparo (Harvard Medical School); strain ET096 | — |
| EST111 | ΔPol Y1 | PY79 *yqjH::loxP spec* | Transformation: *yqjH::loxP spec* → EST003 | This study |
| EST115 | Pol Y1 loxP spec | PY79 *yqjH loxP spec* | Transformation: *yqjH loxP spec* → EST003 | This study |
| EST117 | ΔPol Y2 | PY79 *yqjW::loxP spec* | Transformation: *yqjW::loxP spec* → EST003 | This study |
| EST119 | Pol Y1-CBM-mut1 | PY79 *yqjH-CBM-mut1 loxP spec* | Transformation: *yqjH-CBM-mut1 loxP spec* → EST003 | This study |
| EST121 | Pol Y1-CBM-mut2 | PY79 *yqjH-CBM-mut2 loxP spec* | Transformation: *yqjH-CBM-mut2 loxP spec* → EST003 | This study |
| EST139 | Pol Y1-Dendra2 | PY79 *yqjH-dendra2 loxP spec* | Transformation: *yqjH-dendra2 loxP spec* → EST003 | This study |
| EST141 | Pol Y1-mYPet | PY79 *yqjH-mYPet loxP spec* | Transformation: *yqjH-mYPet loxP spec* → EST003 | This study |
| EST143 | Pol Y1-Halo | PY79 *yqjH-halo loxP spec* | Transformation: *yqjH-halo loxP spec* → EST003 | This study |
| EST191 | Pol Y1-cat-mut | PY79 *yqjH-cat-mut loxP spec* | Transformation: *yqjH-cat-mut loxP spec* → EST003 | This study |
| EST197 | Pol Y1-Halo DnaX-mYPet | PY79 *yqjH-halo loxP spec dnaX-mYpet cat Ω pWX340a* | Transformation: EST143 → EST053 | This study |

| | | | | |
|---|---|---|---|---|
| EST215 | Pol Y1-CBM-mut1-Halo | PY79 *yqjH-CBM-mut1-halo loxP spec* | Transformation: *yqjH-CBM-mut1-halo loxP spec* → EST003 | This study |
| EST217 | Pol Y1-CBM-mut2-Halo | PY79 *yqjH-CBM-mut2-halo loxP spec* | Transformation: *yqjH-CBM-mut2-halo loxP spec* → EST003 | This study |
| EST219 | Pol Y1-CBM-mut1-Halo DnaX-mYPet | PY79 *yqjH-CBM-mut1-halo loxP spec dnaX-mYpet cat Ω pWX340a* | Transformation: EST215 → EST053 | This study |
| EST221 | Pol Y1-CBM-mut2-Halo DnaX-mYPet | PY79 *yqjH-CBM-mut2-halo loxP spec dnaX-mYpet cat Ω pWX340a* | Transformation: EST217 → EST053 | This study |
| EST229 | Pol Y1-cat-mut-Halo | PY79 *yqjH-cat-mut-halo loxP spec* | Transformation: *yqjH-cat-mut-halo loxP spec* → EST003 | This study |
| EST243 | Pol Y1-cat-mut-Halo DnaX-mYPet | PY79 *yqjH-cat-mut-halo loxP spec dnaX-mYpet cat Ω pWX340a* | Transformation: EST229 → EST053 | This study |

### Table S3: Imaging dataset size

| **Dataset/Condition** | **Figure(s)** | **Number of Days** | **Number of Replicates** | **Number of Cells** | **Number of Tracks or Foci** |
|---|---|---|---|---|---|
| DnaX cellular localization<br>Untreated | 3B | 8 | 8 | 816 | 1,363 |
| WT Pol Y1 cellular localization<br>Untreated | 3C, S1A | 8 | 8 | 816 | 26,602 |
| DnaX cellular localization<br>10 μM 4-NQO (1 h) | 3D | 4 | 6 | 760 | 1,522 |
| WT Pol Y1 cellular localization<br>10 μM 4-NQO (1 h) | 3E, S1B | 4 | 6 | 760 | 40,469 |
| WT Pol Y1-DnaX $g(r)$<br>Untreated (All trajectories) | 4A, 4C, S4C | 8 | 8 | 816 | 25,009 |
| WT Pol Y1-DnaX $g(r)$<br>Untreated (Static: $D^* < 0.14$ μm$^2$/s) | 4A | 8 | 8 | 816 | 8,465 |
| WT Pol Y1-DnaX $g(r)$<br>Untreated (Mobile: $D^* > 0.14$ μm$^2$/s) | 4A | 8 | 8 | 816 | 16,544 |
| Pol Y1-CBM-mut1-DnaX $g(r)$<br>Untreated | 4B | 4 | 4 | 636 | 8,298 |
| Pol Y1-CBM-mut2-DnaX $g(r)$<br>Untreated | 4B | 3 | 3 | 872 | 14,694 |
| Pol Y1-cat-mut-DnaX $g(r)$<br>Untreated | 4B | 3 | 3 | 593 | 10,389 |
| WT Pol Y1-DnaX $g(r)$<br>1 μM 4-NQO (1 h) | 4C | 3 | 4 | 621 | 9,357 |
| WT Pol Y1-DnaX $g(r)$<br>10 μM 4-NQO (1 h) | 4C, S4C | 4 | 6 | 760 | 39,558 |
| WT Pol Y1-DnaX $g(r)$<br>10 μM 4-NQO (2 h) | 4C | 2 | 3 | 238 | 16,728 |

| | | | | | |
|---|---|---|---|---|---|
| WT Pol Y1 $D^*$<br>Untreated (All trajectories) | 5A, 5H, S5A | 8 | 8 | 816 | 24,544 |
| WT Pol Y1 $D^*$<br>Untreated (Colocalized: < 200 nm to DnaX) | 5B | 8 | 8 | 816 | 4,856 |
| WT Pol Y1 $D^*$<br>Untreated (Not colocalized: > 200 nm to DnaX-mYPet) | 5C | 8 | 8 | 816 | 18,264 |
| WT Pol Y1 $D^*$<br>10 μM 4-NQO (1 h) | 5D, 5H, S5C | 4 | 6 | 760 | 37,383 |
| Pol Y1-CBM-mut1 $D^*$<br>Untreated | 5E | 4 | 4 | 636 | 8,010 |
| Pol Y1-CBM-mut2 $D^*$<br>Untreated | 5F | 3 | 3 | 872 | 13,953 |
| Pol Y1-cat-mut $D^*$<br>Untreated | 5G | 3 | 3 | 593 | 10,085 |
| WT Pol Y1 $D^*$<br>Fixed | S1B | 2 | 2 | 282 | 3,364 |
| WT Pol Y1 Mean PSF Width<br>Fixed (Long exposure) | S1C | 2 | 2 | 294 | 2,358 |
| False positive spots cellular localization<br>Untreated | S1A | 2 | 2 | 158 | 113 |
| False positive spots cellular localization<br>10 μM 4-NQO (1 h) | S1B | 2 | 2 | 268 | 310 |
| DnaX cellular localization<br>Untreated (Long exposure) | S1C | 8 | 8 | 916 | 1,544 |
| DnaX cellular localization<br>10 μM 4-NQO (1 h) (Long exposure) | S1D | 4 | 6 | 820 | 1,668 |
| WT Pol Y1 cellular localization<br>Untreated (Long exposure) | S1E | 8 | 8 | 916 | 19,538 |
| WT Pol Y1 cellular localization<br>10 μM 4-NQO (1 h) (Long exposure) | S1F | 4 | 6 | 820 | 21,905 |
| WT Pol Y1 cellular localization<br>1 μM 4-NQO (1 h) | S1G | 3 | 4 | 621 | 9,732 |
| WT Pol Y1 cellular localization<br>10 μM 4-NQO (2 h) | S1H | 2 | 3 | 238 | 17,142 |
| Pol Y1-CBM-mut1 cellular localization<br>Untreated | S3A | 4 | 4 | 636 | 8,933 |
| Pol Y1-CBM-mut1 cellular localization<br>10 μM 4-NQO (1 h) | S3B | 2 | 3 | 600 | 18,634 |
| Pol Y1-CBM-mut2 cellular localization<br>Untreated | S3C | 3 | 3 | 872 | 15,382 |
| Pol Y1-CBM-mut2 cellular localization<br>10 μM 4-NQO (1 h) | S3D | 2 | 3 | 697 | 21,630 |
| Pol Y1-cat-mut cellular localization<br>Untreated | S3E | 3 | 3 | 593 | 10,944 |
| Pol Y1-cat-mut cellular localization<br>10 μM 4-NQO (1 h) | S3F | 3 | 3 | 674 | 24,904 |
| WT Pol Y1-DnaX $g(r)$<br>Untreated (Long exposure) | S4B | 8 | 8 | 916 | 18,545 |
| WT Pol Y1-DnaX $g(r)$<br>10 μM 4-NQO (1 h) (Long exposure) | S4B | 4 | 6 | 820 | 21,249 |
| Pol Y1-CBM-mut1-DnaX $g(r)$<br>10 μM 4-NQO (1 h) | S4C | 2 | 3 | 600 | 18,170 |

| | | | | | |
|---|---|---|---|---|---|
| Pol Y1-CBM-mut2-DnaX *g*(*r*)<br>10 μM 4-NQO (1 h) | S4C | 2 | 3 | 697 | 20,573 |
| Pol Y1-cat-mut-DnaX *g*(*r*)<br>10 μM 4-NQO (1 h) | S4C | 3 | 3 | 674 | 24,296 |
| False positive spots $D^*$<br>Untreated | S5B | 2 | 2 | 158 | 98 |
| False positive spots $D^*$<br>10 μM 4-NQO (1 h) | S5D | 2 | 2 | 268 | 277 |
| WT Pol Y1 $D^*$<br>1 μM 4-NQO (1 h) | S5E | 3 | 4 | 621 | 9,029 |
| WT Pol Y1 $D^*$<br>10 μM 4-NQO (2 h) | S5F | 2 | 3 | 238 | 15,976 |
| Pol Y1-CBM-mut1 $D^*$<br>10 μM 4-NQO (1 h) | S5G | 2 | 3 | 600 | 16,892 |
| Pol Y1-CBM-mut2 $D^*$<br>10 μM 4-NQO (1 h) | S5H | 2 | 3 | 697 | 19,615 |
| Pol Y1-cat-mut $D^*$<br>10 μM 4-NQO (1 h) | S5I | 3 | 3 | 674 | 23,101 |
| WT Pol Y1 binding lifetime<br>Untreated | S6B,<br>S6C,<br>S6E | 8 | 8 | 916 | 19,538 |
| WT Pol Y1 binding lifetime<br>Fixed | S6B,<br>S6C,<br>S6D | 2 | 2 | 294 | 6,207 |
| Pol Y1-cat-mut binding lifetime<br>Untreated | S6B, S6F | 3 | 3 | 645 | 13,235 |
| WT Pol Y1 binding lifetime<br>10 μM 4-NQO (1 h) | S6C,<br>S6G | 4 | 6 | 820 | 21,905 |

**Table S4: Value of the mean radial distribution function *g*(*r*) for Pol Y1-DnaX colocalization at the second smallest value of *r* (generally the maximum of the *g*(*r*) curve) and the standard error of the mean (S.E.M.) at that *r* value for the 100 calculated *g*(*r*) curves**

| Figure(s) | Protein | Condition | *g*(*r*) ± S.E.M. |
|---|---|---|---|
| 4A, 4C, S4C | WT Pol Y1 | Untreated (All trajectories) | 2.84 ± 0.01 |
| 4A | WT Pol Y1 | Untreated (Static: $D^* < 0.14$ μm$^2$/s) | 5.95 ± 0.04 |
| | WT Pol Y1 | Untreated (Mobile: $D^* > 0.14$ μm$^2$/s) | 1.112 ± 0.006 |
| 4B | Pol Y1-CBM-mut1 | Untreated | 0.157 ± 0.001 |
| | Pol Y1-CBM-mut2 | Untreated | 0.0938 ± 0.0005 |
| | Pol Y1-cat-mut | Untreated | 3.20 ± 0.02 |
| 4C | WT Pol Y1 | 1 μM 4-NQO (1 h) | 2.14 ± 0.02 |
| | WT Pol Y1 | 10 μM 4-NQO (1 h) | 2.370 ± 0.007 |
| | WT Pol Y1 | 10 μM 4-NQO (2 h) | 2.54 ± 0.01 |
| S4B | WT Pol Y1 | Untreated; Long exposure | 2.76 ± 0.01 |
| | WT Pol Y1 | 10 μM 4-NQO (1 h); Long exposure | 2.79 ± 0.01 |
| S4C | Pol Y1-CBM-mut1 | 10 μM 4-NQO (1 h) | 0.536 ± 0.003 |
| | Pol Y1-CBM-mut2 | 10 μM 4-NQO (1 h) | 0.435 ± 0.002 |
| | Pol Y1-cat-mut | 10 μM 4-NQO (1 h) | 3.05 ± 0.01 |

**Table S5: Pol Y1-Halo diffusion coefficient distribution fit parameters (± uncertainties from 95% fit confidence intervals)**

| Strain/Condition | $D_1$ (μm²/s) | $A_1$ | $D_2$ (μm²/s) | $A_2$ | $D_3$ (μm²/s) | $A_3$ |
|---|---|---|---|---|---|---|
| WT Pol Y1 Untreated (All) | 0.080 ± 0.005 | 0.281 ± 0.028 | 0.976 ± 0.077 | 0.468 ± 0.030 | 0.234 ± 0.030 | 0.250 ± 0.058 |
| WT Pol Y1 Untreated (< 200 nm to DnaX-mYPet) | 0.074 ± 0.006 | 0.440 ± 0.069 | 0.734 ± 0.187 | 0.209 ± 0.046 | 0.186 ± 0.032 | 0.351 ± 0.256 |
| WT Pol Y1 Untreated (> 200 nm to DnaX-mYPet) | 0.084 ± 0.005 | 0.243 ± 0.0234 | 0.996 ± 0.065 | 0.522 ± 0.028 | 0.253 ± 0.031 | 0.235 ± 0.051 |
| WT Pol Y1 10 μM 4-NQO (1 h) | 0.079 ± 0.004 | 0.282 ± 0.026 | 1.022 ± 0.092 | 0.459 ± 0.034 | 0.253 ± 0.032 | 0.259 ± 0.060 |
| Pol Y1-CBM-mut1 Untreated | 0.081 ± 0.005 | 0.108 ± 0.010 | 1.117 ± 0.041 | 0.682 ± 0.023 | 0.325 ± 0.028 | 0.210 ± 0.033 |
| Pol Y1-CBM-mut2 Untreated | 0.086 ± 0.005 | 0.086 ± 0.008 | 1.159 ± 0.031 | 0.725 ± 0.018 | 0.344 ± 0.025 | 0.188 ± 0.026 |
| Pol Y1-cat-mut Untreated | 0.079 ± 0.005 | 0.336 ± 0.038 | 1.009 ± 0.085 | 0.427 ± 0.026 | 0.204 ± 0.029 | 0.237 ± 0.064 |
| WT Pol Y1 1 μM 4-NQO (1 h) | 0.079 ± 0.005 | 0.314 ± 0.033 | 1.047 ± 0.085 | 0.474 ± 0.028 | 0.220 ± 0.034 | 0.213 ± 0.061 |
| WT Pol Y1 10 μM 4-NQO (2 h) | 0.081 ± 0.004 | 0.280 ± 0.027 | 0.994 ± 0.072 | 0.461 ± 0.026 | 0.230 ± 0.026 | 0.260 ± 0.053 |
| Pol Y1-CBM-mut1 10 μM 4-NQO (1 h) | 0.084 ± 0.005 | 0.118 ± 0.010 | 1.136 ± 0.041 | 0.687 ± 0.024 | 0.342 ± 0.032 | 0.195 ± 0.034 |
| Pol Y1-CBM-mut2 10 μM 4-NQO (1 h) | 0.076 ± 0.005 | 0.110 ± 0.009 | 1.128 ± 0.039 | 0.722 ± 0.023 | 0.332 ± 0.035 | 0.168 ± 0.032 |
| Pol Y1-cat-mut 10 μM 4-NQO (1 h) | 0.077 ± 0.005 | 0.357 ± 0.042 | 1.053 ± 0.1197 | 0.423 ± 0.034 | 0.212 ± 0.039 | 0.220 ± 0.076 |

**Table S6: Pol Y1-Halo diffusion coefficient distribution fit parameters (± uncertainties from 95% fit confidence intervals)**

| Strain/Condition | $\tau_{bleach}$ (s) | $\tau_{app}$ (s) | $\tau_{bound}$ (s) |
|---|---|---|---|
| WT Pol Y1 Fixed | 1.10 ± 0.05 | — | — |
| WT Pol Y1 Untreated | — | 0.70 ± 0.02 | 1.9 ± 0.4 |
| Pol Y1-cat-mut Untreated | — | 0.92 ± 0.03 | 6 ± 3 |
| WT Pol Y1 10 μM 4-NQO (1 h) | — | 0.73 ± 0.02 | 2.2 ± 0.5 |

**Table S7: Fold change in number of colony forming units per mL (CFUs/mL) for imaging cultures after different treatments (± standard deviation)**

| **Condition** | **Untreated** | **DMF Only** | **1 μM 4-NQO (1 h)** | **10 μM 4-NQO (1 h)** | **10 μM 4-NQO (2 h)** | **50 μM 4-NQO (1 h)** |
|---|---|---|---|---|---|---|
| **Fold Change in CFUs/mL** | 2.09 ± 0.08 | 2.3 ± 0.6 | 2.1 ± 0.2 | 0.7 ± 0.2 | 1.5 ± 0.3 | 0.06 ± 0.02 |

### Supplementary Methods

*Overview of strain construction strategy:*

Bacterial strains containing fluorescent protein fusions or other modifications were constructed by transformation of either double-stranded DNA (dsDNA) fragments or genomic DNA bearing 1 – 2 kilobase (kb) homology arms for incorporation into the chromosome. dsDNA fragments containing desired modifications were synthesized by polymerase chain reaction (PCR) amplification and Gibson assembly.(4) Genomic DNA was extracted with phenol-chloroform and ethanol precipitation, resuspended in Tris-EDTA (TE) buffer, and transformed without further purification. Recipient strains were grown at 37 °C in BMK Complete medium (60 mM $K_2HPO_4$, 38 mM $KH_2PO_4$, 111 mM D-glucose, 3 mM sodium citrate, 0.0022% ferric ammonium citrate, 15 mM L-aspartic acid potassium salt, 10 mM $MgSO_4$, 0.05% yeast extract) to induce competence. Transformants were selected on LB Lennox agar plates containing the appropriate antibiotic, followed by one round of streak purification after the initial selection step. Newly transformed modifications were validated by PCR amplification of genomic DNA followed by Sanger DNA sequencing. Preexisting modifications were checked by streaking on antibiotic plates. Antibiotic concentrations used were 100 μg/mL spectinomycin and 5 μg/mL chloramphenicol. Tables S1 and S2 list all oligonucleotides and bacterial strains used in this study. Construction details for all new strains are summarized below.

The Pol Y1-Halo construct for microscopy was designed as a C-terminal fusion to the self-labeling HaloTag(5) with an 11 amino acid linker (GSGQGSGPGSG) between the Pol Y1 C-terminus and the HaloTag N-terminus. Similar C-terminal Pol Y1 fusions were made to the monomeric YFP variant mYPet(6) (using the 8 amino acid linker LEGSGQGP) and the photoconvertible green fluorescent protein (GFP) variant Dendra2(7, 8) (using the 8 amino acid linker LEGSGGSG). The DnaX replisome marker contained a C-terminal fusion to mYPet with an 8 amino acid linker (LEGSGQGP). The Pol Y1 catalytically inactive mutant contained the D108A E109A mutation to the catalytic residues in the polymerase active site.(9) The Pol Y1 CBM-mut1 and CBM-mut2 mutants contained mutations (QADAF and ALDLA respectively) to the WT clamp-binding motif sequence (QLDLF).

The *yqjW* gene is the first gene in an operon.(10) To minimize effects on the expression of the downstream gene, *yqjX*, the N-terminal 18 base pairs (bp) and the C-terminal 24 bp of *yqjW* (including the stop codon) were retained in the *yqjW::spec* knockout strain.

*Detailed strain construction information:*

**EST111:** ΔPol Y1. The *yqjH* upstream was amplified from strain EST003 using oligonucleotides oEST037 and oEST038. The loxP spec cassette was amplified from strain EST081 using oligonucleotides oEST039 and oEST040. The *yqjH* downstream was amplified from strain

EST003 using oligonucleotides oEST041 and oEST028. The three fragments were joined by Gibson assembly and transformed into strain EST003.

**EST115:** Pol Y1 loxP spec. The *yqjH* gene and upstream were amplified from strain EST003 using oligonucleotides oEST023 and oEST070. The loxP spec cassette was amplified from strain EST081 using oligonucleotides oEST032 and oEST008. The *yqjH* downstream was amplified from strain EST003 using oligonucleotides oEST071 and oEST028. The three fragments were joined by Gibson assembly and transformed into strain EST003.

**EST117:** ΔPol Y2. The *yqjW* upstream was amplified from strain EST003 using oligonucleotides oEST049 and oEST072. The loxP spec cassette was amplified from strain EST081 using oligonucleotides oEST032 and oEST008. The *yqjW* downstream was amplified from strain EST003 using oligonucleotides oEST073 and oEST048. The three fragments were joined by Gibson assembly and transformed into strain EST003.

**EST119:** Pol Y1-CBM-mut1. The N-terminal portion of *yqjH* (from the N-terminus to the clamp-binding motif) and the upstream were amplified from strain EST115 using oligonucleotides oEST023 and oEST058. The C-terminal portion of *yqjH* (from the clamp-binding motif to the C-terminus) and the downstream were amplified from strain EST115 using oligonucleotides oEST059 and oEST028. Oligonucleotides oEST058 and oEST059 contained the CBM-mut1 mutations. The two fragments were joined by Gibson assembly and transformed into strain EST003.

**EST121:** Pol Y1-CBM-mut2. The N-terminal portion of *yqjH* (from the N-terminus to the clamp-binding motif) and the upstream were amplified from strain EST115 using oligonucleotides oEST023 and oEST060. The C-terminal portion of *yqjH* (from the clamp-binding motif to the C-terminus) and the downstream were amplified from strain EST115 using oligonucleotides oEST061 and oEST028. Oligonucleotides oEST060 and oEST061 contained the CBM-mut2 mutations. The two fragments were joined by Gibson assembly and transformed into strain EST003.

**EST139:** Pol Y1-Dendra2. The *yqjH* gene and upstream were amplified from strain EST003 using oligonucleotides oEST023 and oEST029. The linker and Dendra2 were amplified from strain EST081 using oligonucleotides oEST033 and oEST034. The loxP spec cassette and *yqjH* downstream were amplified from strain EST115 using oligonucleotides oEST032 and oEST028. The three fragments were joined by Gibson assembly and transformed into strain EST003.

**EST141:** Pol Y1-mYPet. The *yqjH* gene and upstream were amplified from strain EST003 using oligonucleotides oEST023 and oEST029. The linker and mYPet were amplified from strain EST057 using oligonucleotides oEST035 and oEST036. The loxP spec cassette and *yqjH* downstream were amplified from strain EST115 using oligonucleotides oEST032 and oEST028. The three fragments were joined by Gibson assembly and transformed into strain EST003.

**EST143:** Pol Y1-Halo. The *yqjH* gene and upstream were amplified from strain EST003 using oligonucleotides oEST023 and oEST029. The linker and HaloTag were amplified from strain EST083 using oligonucleotides oEST030 and oEST031. The loxP spec cassette and *yqjH* downstream were amplified from strain EST115 using oligonucleotides oEST032 and oEST028. The three fragments were joined by Gibson assembly and transformed into strain EST003.

**EST191:** Pol Y1-cat-mut. The N-terminal portion of *yqjH* (from the N-terminus to the catalytic site) and the upstream were amplified from strain EST115 using oligonucleotides oEST037 and oEST062. The C-terminal portion of *yqjH* (from the clamp-binding motif to the C-terminus) and the downstream were amplified from strain EST115 using oligonucleotides oEST063 and oEST028. Oligonucleotides oEST062 and oEST063 contained the catalytic site mutations. The two fragments were joined by Gibson assembly and transformed into strain EST003.

**EST197:** Pol Y1-Halo DnaX-mYPet. The *yqjH-halo loxP spec* allele was transferred from strain EST143 to strain EST053 by transformation with genomic DNA.

**EST215:** Pol Y1-CBM-mut1-Halo. The *yqjH* gene (containing the CBM-mut1 mutation) and upstream were amplified from strain EST119 using oligonucleotides oEST023 and oEST029. The linker and HaloTag were amplified from strain EST083 using oligonucleotides oEST030 and oEST031. The loxP spec cassette and *yqjH* downstream were amplified from strain EST115 using oligonucleotides oEST032 and oEST028. The three fragments were joined by Gibson assembly and transformed into strain EST003.

**EST217:** Pol Y1-CBM-mut2-Halo. The *yqjH* gene (containing the CBM-mut2 mutation) and upstream were amplified from strain EST121 using oligonucleotides oEST023 and oEST029. The linker and HaloTag were amplified from strain EST083 using oligonucleotides oEST030 and oEST031. The loxP spec cassette and *yqjH* downstream were amplified from strain EST115 using oligonucleotides oEST032 and oEST028. The three fragments were joined by Gibson assembly and transformed into strain EST003.

**EST219:** Pol Y1-CBM-mut1-Halo DnaX-mYPet. The *yqjH-CBM-mut1-halo loxP spec* allele was transferred from strain EST215 to strain EST053 by transformation with genomic DNA.

**EST221:** Pol Y1-CBM-mut2-Halo DnaX-mYPet. The *yqjH-CBM-mut2-halo loxP spec* allele was transferred from strain EST217 to strain EST053 by transformation with genomic DNA.

**EST229:** Pol Y1-cat-mut-Halo. The *yqjH* gene (containing the catalytic site mutations) and upstream were amplified from strain EST191 using oligonucleotides oEST023 and oEST029. The linker and HaloTag were amplified from strain EST083 using oligonucleotides oEST030 and oEST031. The loxP spec cassette and *yqjH* downstream were amplified from strain EST115 using oligonucleotides oEST032 and oEST028. The three fragments were joined by Gibson assembly and transformed into strain EST003.

**EST243:** Pol Y1-cat-mut-Halo DnaX-mYPet. The *yqjH-cat-mut-halo loxP spec* allele was transferred from strain EST229 to strain EST053 by transformation with genomic DNA.
